# CBFA2T3–GLIS2 establishes transcriptionally permissive enhancer hypermethylation associated with venetoclax response in pediatric AML

**DOI:** 10.64898/2026.10.01.756000

**Authors:** Arundhati Chavan, Rhonda Ries, Pritam Biswas, Jason E Farrar, Soheil Meshinchi, Samrat Roy Choudhury

## Abstract

CBFA2T3-GLIS2 (C/G) defines a high-risk pediatric AML subtype in which *de novo* enhancer rewiring sustains an aggressive leukemic transcriptional state. The early epigenomic mechanisms reinforcing this program and linking it to leukemic survival remain incompletely defined. Here, we identify enhancer-proximal hypermethylation as a noncanonical, transcription-permissive feature of C/G AML. Using primary C/G-positive AML samples and a developmentally relevant C/G-transformed cord blood hematopoietic stem/progenitor cell model, we integrated DNA methylation, chromatin state, fusion occupancy, and gene expression across early and leukemia-stage contexts. Early C/G-associated hypermethylated regions localized near H3K27ac/H3K4me1-marked active or primed cis-regulatory elements, were enriched for ERG/ETS-like motifs, and showed related architecture in leukemia-stage contexts. *DNMT3B* was prominently increased in C/G contexts, and its loss selectively altered methylation-associated gene expression without measurably changing global 5mC. Genetic DNMT3B loss increased venetoclax sensitivity in vitro, while pharmacologic DNMT3B inhibition with Nanaomycin-A similarly enhanced venetoclax response. During venetoclax treatment, DNMT3B-deficient xenografts showed reduced disease burden and prolonged survival. To assess whether this interaction extended to clinically used hypomethylating agents, azacitidine or decitabine plus venetoclax induced apoptosis-related transcriptional programs and mitochondrial depolarization. These findings identify hypermethylated active cis-regulatory elements as C/G-associated states linked to transcriptional dysregulation and venetoclax response.

## Introduction

Pediatric acute myeloid leukemia (pAML) is often driven by chromosomal fusions that alter transcriptional regulation during hematopoietic development. The CBFA2T3–GLIS2 (C/G) fusion defines a highly aggressive pAML subtype, enriched in acute megakaryoblastic leukemia (FAB M7) and predominantly observed in infants and young children, but also detected across non-M7 lineage contexts [1, 2]. Clinically, C/G⁺ patients frequently exhibit extramedullary disease, high residual burden, early relapse after intensive therapy or transplantation, and poor survival, placing this fusion among the worst-prognosis pAML subtypes [3-7].

Mechanistic studies have shown that C/G redirects transcriptional complexes to atypical cis-regulatory elements (CREs) and generates *de novo* enhancer and super-enhancer states that sustain leukemic transcription [8, 9]. Human cord blood progenitor systems have further shown that C/G can initiate a patient-like leukemic transcriptional program in a developmentally relevant hematopoietic context [5, 6]. These studies establish C/G as an enhancer-rewiring oncogene and provide a tractable window to interrogate early fusion-instigated epigenomic remodeling. However, how C/G-induced enhancer states become coupled to DNA methylation (DNAm), how these states evolve from early fusion expression to leukemia-stage regulatory programs, and whether they influence leukemic survival remain incompletely defined.

Our previous work identified a noncanonical DNAm architecture in C/G⁺ pAML, in which hypermethylated CREs can remain transcriptionally permissive and associate with active gene expression [10]. This raised a specific question: whether enhancer-proximal hypermethylation emerges early after C/G expression and contributes to active regulatory states. Because prior occupancy data place C/G at the *DNMT3B* promoter, *DNMT3B* is prominently increased in C/G⁺ AML, and CRE hypermethylation is recurrent, we tested whether *DNMT3B* contributes to selected methylation-linked regulatory states. We further asked whether DNMT3B loss is accompanied by changes in DNMT1-UHRF1 maintenance machinery that could help explain preserved bulk 5mC despite locus-specific regulatory effects [5, 10].

Using a developmentally relevant umbilical cord blood-derived hematopoietic stem/progenitor cell (UCB-HSPC) model, we profile early C/G-associated DNAm and chromatin remodeling, define motif architecture within hypermethylated active/primed CRE neighborhoods, and compare selected regulatory states with leukemia-stage C/G⁺ contexts. Functionally, DNMT perturbation increased venetoclax sensitivity and was accompanied by apoptosis-related transcriptional and mitochondrial changes. Together, these findings identify enhancer-proximal hypermethylation as an early, transcriptionally permissive regulatory feature in high-risk C/G-driven pAML.

## Materials and Methods

### Patient samples and cell lines

Primary pAML samples from COG trials AAML0531, AAML1031, and AAML03P1 were compared with NBM from healthy donors <18 years, as previously described [10]. The study included C/G^+^ AML cell lines M07e and WSU-AML, C/G− controls MV4-11 and AML-193, and reference UCB-CD34^+^ HSPCs and megakaryocytic (MK) progenitors where indicated. Cohort composition, dataset accessions, culture conditions, cell authentication, and flow cytometry are provided in Supplementary Information (SI).

### C/G-transformed cord blood model

C/G–GFP- and GFP-transduced UCB-CD34^+^ cells were generated by lentiviral transduction and endothelial co-culture as previously described [5, 6]. Day 42 cultures were harvested after acquisition of the C/G-associated FOLR1⁺ phenotype and profiled by flow cytometry, RNA-seq, CUT&RUN for four histone marks (H3K4me1, H3K27ac, H3K4me3, and H3K27me3), and methyl-enrichment enzymatic methyl sequencing (meEM-seq). Transduction, endothelial co-culture, FACS sorting, and assay-specific workflows are provided in SI.

### DNAm, chromatin, and fusion-occupancy integration

Published patient EPIC methylation data and M07e CBFA2T3/GLIS2 CUT&RUN maps were used to anchor C/G-associated DNAm changes to leukemia-stage fusion occupancy [10]. Early fusion-induced regulatory changes were captured by profiling FOLR1⁺ C/G-transformed UCB cells and GFP controls by meEM-seq and CUT&RUN for H3K4me1, H3K27ac, H3K4me3, and H3K27me3. Regulatory integration focused on H3K4me1/H3K27ac-defined active/primed chromatin, with H3K4me3/H3K27me3 used to assess promoter-associated and repressive chromatin redistribution. UCB DMRs, patient DMCs, chromatin profiles, and leukemia-stage C/G occupancy were integrated to define methylation-linked regulatory states across early developmental and leukemia-stage contexts. DMC/DMR calling, peak processing, coordinate conversion, patient DMC clustering, motif proximity analysis, and genome-browser visualization are provided in SI.

### Transcriptomic and regulatory enrichment analyses

Differential expression was analyzed for C/G⁺ pAML versus NBM and C/G-transformed versus GFP-transduced UCB-CD34⁺ cells, followed by concordance analysis between patient and UCB datasets. Patient- and UCB-ranked transcriptomes were used for preranked GSEA, and shared C/G-associated genes were tested for MSigDB Hallmark enrichment. Active/primed hypermethylated regions from the UCB model and patient DMR-like CpG clusters in the leukemia-stage M07e chromatin context were analyzed for motif enrichment and ERG/ETS-like motif proximity. RNA-seq from DNMT3B-KO M07e cells and AZA-, VEN-, and AZA+VEN-treated M07e cells was used to assess apoptosis-related transcriptional responses.

### DNMT3B perturbation, rescue, and therapeutic response assays

DNMT3B KO and gRNA-resistant DNMT3B rescue were performed in M07e cells, followed by immunoblot validation, global 5mC quantification by LC-MS, and locus-specific DNAm/expression analyses at selected C/G-linked regulatory targets. VEN response was tested after genetic DNMT3B loss and Nan-A-mediated DNMT3B-biased perturbation. AZA and DAC were evaluated with VEN in concurrent-combination and HMA-priming schedules. Cell viability was measured across dose ranges, with IC50 and AUC calculated from biological replicates. In vivo VEN response was tested in M07e WT and DNMT3B-KO xenografts using bioluminescence and survival as primary endpoints, with endpoint leukemia burden assessed where indicated. TMRM flow cytometry was used where indicated to assess mitochondrial membrane potential. Perturbation design, rescue validation, methylation/expression assays, drug schedules, xenograft procedures, and TMRM analysis are provided in SI.

### Statistical analysis

Statistical tests were two-sided unless otherwise stated, with false discovery rate correction where appropriate. Data are shown as mean ± SD or SEM from independent biological replicates, as indicated in figure legends. Figure-specific statistical tests and replicate definitions are provided in SI.

## Results

### C/G-positive pAML shows DNMT3B upregulation and promoter-biased overlap between fusion occupancy and hypermethylation

C/G was detected across FAB-defined categories within the TARGET pAML cohort. Among C/G⁺ cases, 43% were classified as M7/acute megakaryoblastic leukemia and 57% occurred outside M7, supporting lineage-diverse distribution of the fusion (Fig. 1A). Consistent with prior clinical reports, C/G⁺ pAML showed the poorest survival among major molecular pAML subgroups; among 36 evaluable C/G⁺ cases with available survival data, median overall survival was 17.2 months and 5-year overall survival was 23.5% (log-rank P < 0.001; Fig. 1B; Table S1). Immunoblotting confirmed endogenous C/G fusion protein expression in M07e and WSU-AML cells, but not in C/G⁻ controls MV4-11 and AML-193. Full-length CBFA2T3 signal was retained in C/G⁺ cells, consistent with the heterozygous inversion structure of the fusion and supporting the use of M07e and WSU-AML as endogenous C/G⁺ models (Fig. 1C).

**Figure 1.**
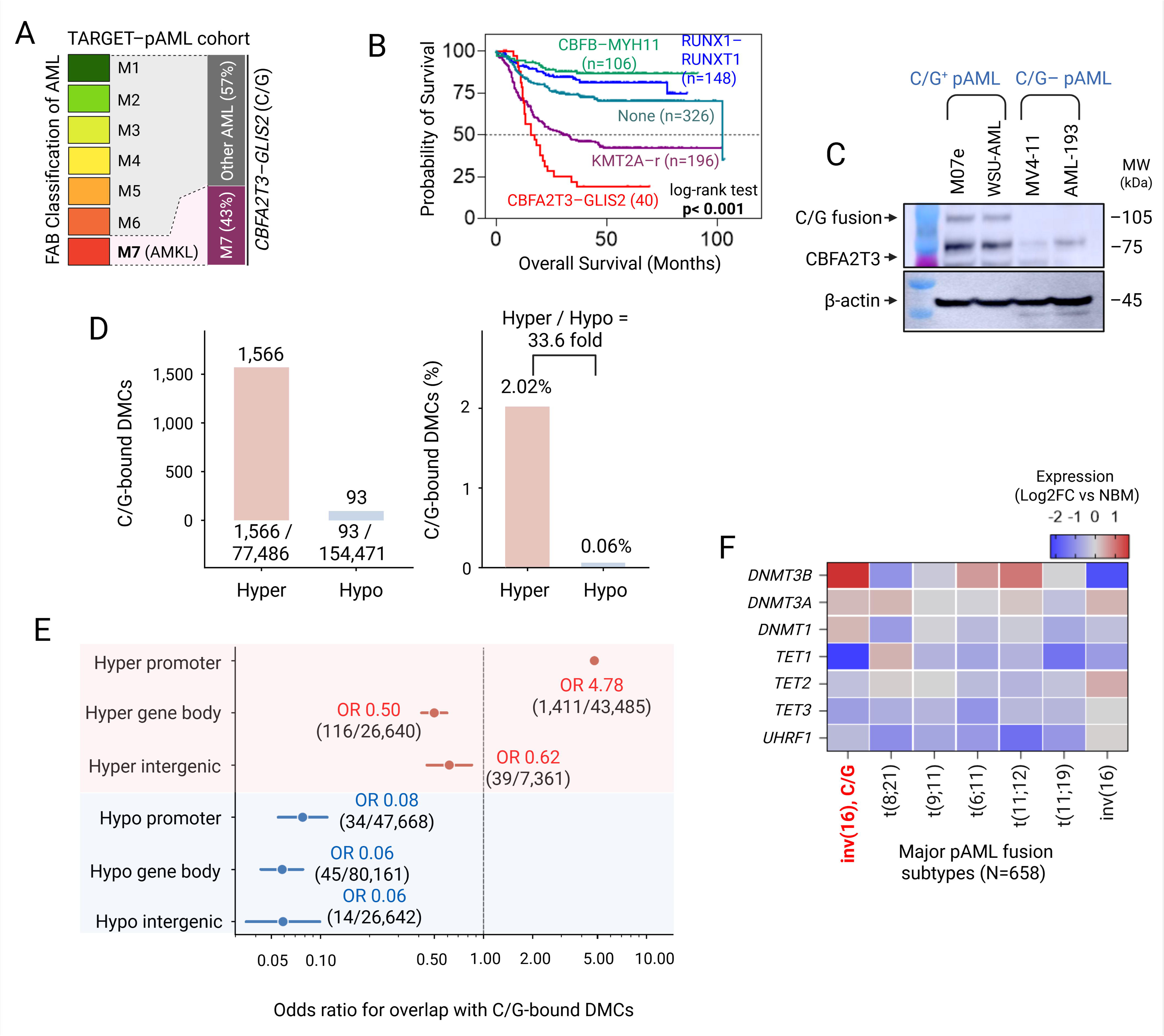
C/G-positive pAML shows DNMT3B upregulation and promoter-biased overlap between fusion occupancy and hypermethylation. (A) Distribution of CBFA2T3–GLIS2 (C/G)⁺ cases across FAB-defined categories within the TARGET pAML cohort. C/G⁺ cases were detected across FAB categories, with 43% classified as M7/acute megakaryoblastic leukemia and 57% occurring outside M7. (B) Kaplan-Meier analysis of overall survival across major molecular pAML subgroups, including CBFB–MYH11, RUNX1–RUNX1T1, KMT2A-rearranged AML, AML without the indicated major fusions (“None”), and C/G⁺ AML. The TARGET cohort included 40 C/G⁺ cases, of which 36 had evaluable survival data and were included in survival analysis. Statistical significance was determined by log-rank test. (C) Immunoblot detection of endogenous C/G fusion protein in C/G⁺ AML cell lines M07e and WSU-AML and C/G⁻ controls MV4-11 and AML-193 using an antibody recognizing the N-terminal CBFA2T3 portion of the fusion. β-actin was used as loading control. (D) Overlap of high-confidence M07e C/G-bound regions with patient differentially methylated CpGs (DMCs) from C/G⁺ pAML versus normal bone marrow. Patient DMCs were defined from EPIC methylation profiling using |Δβ| ≥ 0.05. Among hypermethylated DMCs, 1,566 of 77,486 overlapped C/G-bound regions (2.02%; OR = 2.89; P = 1.36 × 10⁻²³³), compared with 93 of 154,471 hypomethylated DMCs (0.06%; OR = 0.055). (E) Odds-ratio analysis of C/G-bound DMC enrichment stratified by methylation direction and genomic compartment. C/G-bound regions were selectively enriched at promoter-associated hypermethylated CpGs (OR = 4.78; 1,411/43,485; 95% CI, 4.53–5.05), whereas hypermethylated gene body and intergenic CpGs showed odds ratios below unity. Hypomethylated CpGs were depleted from C/G-bound regions across promoter, gene body, and intergenic compartments. (F) Heatmap of DNA methylation and demethylation regulator expression across major pAML fusion-defined subtypes, displayed as log2 fold-change relative to normal bone marrow. DNMT3B showed prominent upregulation in C/G⁺ pAML within the broader DNMT/TET/UHRF1 regulatory landscape. AMKL, acute megakaryoblastic leukemia; C/G, CBFA2T3–GLIS2; CI, confidence interval; DMC, differentially methylated CpG; DNAm, DNA methylation; FAB, French-American-British; NBM, normal bone marrow; OR, odds ratio; OS, overall survival; pAML, pediatric acute myeloid leukemia; TARGET, Therapeutically Applicable Research to Generate Effective Treatments.

We next examined how leukemia-stage fusion occupancy relates to the broader DNAm landscape in primary C/G⁺ pAML. Integration of high-confidence M07e C/G CUT&RUN peaks with patient EPIC methylation data showed that C/G-bound regions overlapped a small but directionally biased subset of patient DMCs. C/G-bound regions overlapped 1,566 of 77,486 hypermethylated DMCs (2.02%; OR = 2.89; P = 1.36 × 10⁻²³³), but only 93 of 154,471 hypomethylated DMCs (0.06%; OR = 0.055; Fig. 1D; Table S2A). Thus, leukemia-stage C/G-bound regions were preferentially associated with hypermethylated rather than hypomethylated CpGs, while accounting for only a focused fraction of total methylation remodeling.

Stratification by genomic annotation showed that this association was concentrated at promoter-associated hypermethylation. C/G-bound regions overlapped 1,411 of 43,485 hypermethylated promoter CpGs (3.24%; OR = 4.78; 95% CI, 4.53–5.05), whereas hypermethylated gene body and intergenic CpGs showed odds ratios below unity (OR = 0.50 and 0.62, respectively; Fig. 1E; Table S2B). Hypomethylated CpGs were depleted from C/G-bound regions across promoter, gene body, and intergenic compartments. In parallel, expression analysis across major pAML fusion subtypes showed prominent DNMT3B upregulation in C/G⁺ pAML within the broader DNMT/TET/UHRF1 regulatory landscape (Fig. 1F; Table S3). Together with prior DNMT3B-promoter occupancy data [10], these results link C/G to DNMT3B upregulation and a focused promoter-hypermethylated compartment. The limited overlap also indicates that most DNAm remodeling is not explained by direct fusion occupancy and likely involves secondary regulatory mechanisms.

### A developmental UCB-HSPC system captures early patient-conserved C/G transcriptional programs

To define early C/G-driven transcriptional reprogramming in a developmentally relevant hematopoietic context, we used a previously established UCB-CD34⁺ HSPC system in which cells were transduced with C/G–GFP or GFP control and maintained on E4ORF1-HUVEC endothelial co-culture (Fig. 2A) [5, 6]. C/G-transformed cultures expanded during serial culture and were harvested at day 42 for phenotypic and molecular profiling (Fig. 2B). Flow cytometry showed acquisition of a C/G-associated CD34⁺/FOLR1⁺ phenotype in C/G-transformed cultures compared with GFP controls, consistent with immunophenotypic features reported in primary C/G⁺ pAML and supporting the use of this system to study early fusion-induced regulatory changes (Fig. 2C, D) [5, 6].

**Figure 2.**
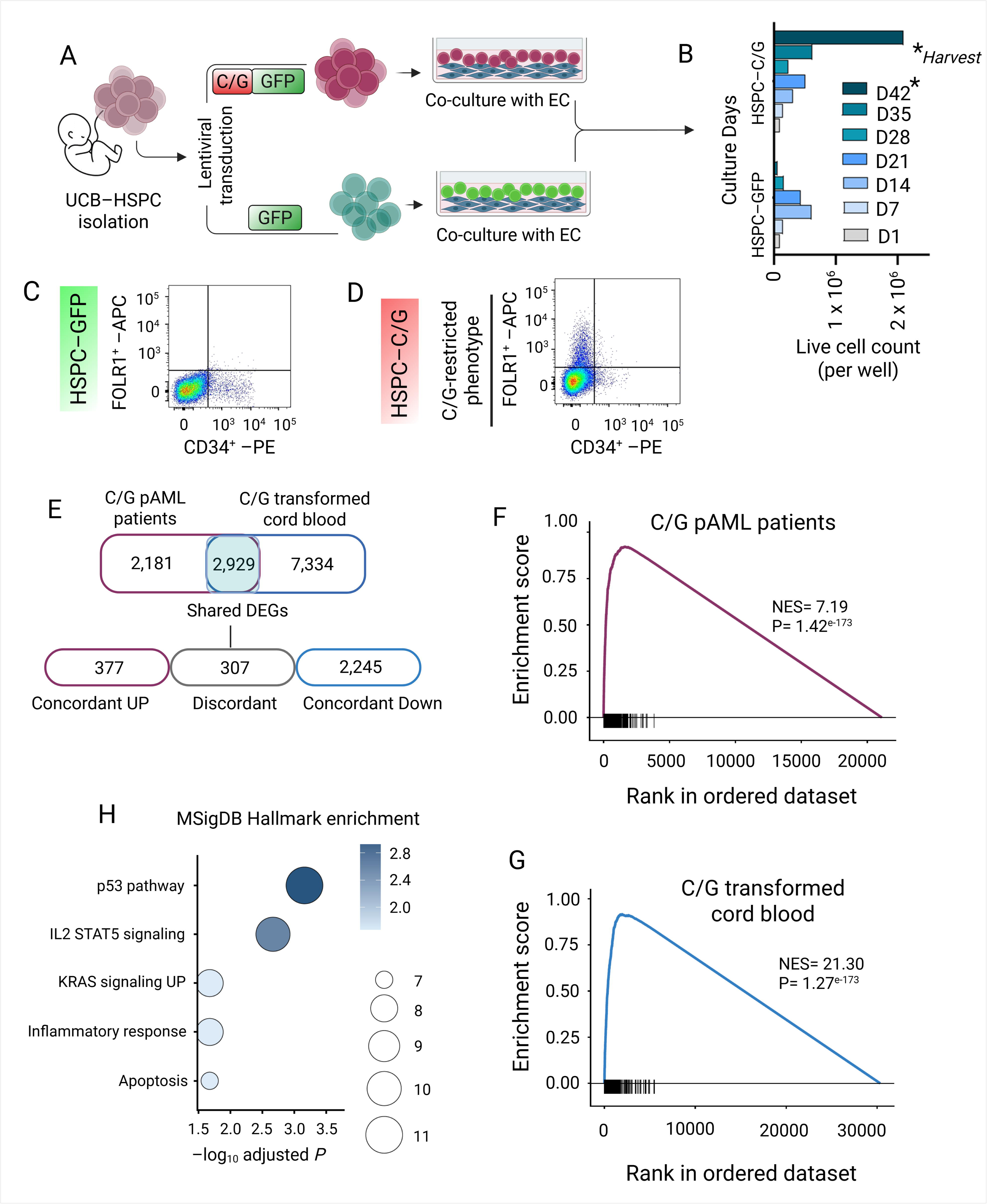
A developmental UCB-HSPC system captures early patient-conserved C/G transcriptional programs. **(A)** Schematic of the C/G-transformed cord blood system. UCB-CD34⁺ HSPCs were transduced with lentiviral CBFA2T3–GLIS2/GFP or GFP control constructs and maintained on E4ORF1-HUVEC endothelial co-culture, as previously described. **(B)** Live-cell expansion of C/G-transformed and GFP-transduced UCB-CD34⁺ cells over serial culture. Cultures were harvested at day 42 for downstream phenotypic and molecular profiling. **(C,D)** Representative flow cytometry plots showing CD34 and FOLR1 expression in GFP-transduced control cells and C/G-transformed cells at day 42. C/G-transformed cells acquired a CD34⁺/FOLR1⁺ phenotype consistent with a C/G-associated pAML immunophenotype. **(E)** Overlap of differentially expressed genes (DEGs) between C/G⁺ pAML versus normal bone marrow and C/G-transformed versus GFP-transduced UCB-CD34⁺ cells. DEGs were defined using |log2FC| ≥ 1.5 and adjusted P < 0.05. The analysis identified 2,929 shared DEGs, 2,181 patient-only DEGs, and 7,334 UCB-only DEGs. Among shared DEGs, 377 were concordantly upregulated, 2,245 were concordantly downregulated, and 307 showed discordant directionality. **(F,G)** Preranked GSEA using the concordantly upregulated shared gene set. The gene set was strongly enriched in both C/G⁺ pAML patient-ranked transcriptomes (NES = 7.19, P = 1.42 × 10⁻¹ ³) and C/G-transformed UCB-ranked transcriptomes (NES = 21.30, P = 1.27 × 10⁻¹ ³). Genes were ranked using a signed significance metric derived from log2FC and P value. **(H)** MSigDB Hallmark enrichment analysis of concordantly upregulated shared genes. Selected enriched programs included p53 pathway, IL2–STAT5 signaling, KRAS signaling, inflammatory response, and apoptosis. Bubble size indicates the number of overlapping genes, and color indicates −log10 adjusted P value. C/G, CBFA2T3–GLIS2; DEG, differentially expressed gene; EC, endothelial co-culture; FOLR1, folate receptor alpha; GFP, green fluorescent protein; GSEA, gene set enrichment analysis; HSPC, hematopoietic stem/progenitor cell; MSigDB, Molecular Signatures Database; NBM, normal bone marrow; NES, normalized enrichment score; pAML, pediatric acute myeloid leukemia; UCB, umbilical cord blood. Figure 2, panel A was created with BioRender.com under publication license JW29XM89G4 (Created in BioRender. Roy choudhury, S. (2026) https://BioRender.com/x80sv29).

We next determined whether this developmental system captured transcriptional programs observed in primary C/G⁺ pAML. Using matched differential-expression thresholds, we compared C/G⁺ pAML versus NBM with C/G-transformed versus GFP-transduced UCB-CD34⁺ cells. This analysis identified 2,929 DEGs shared between patient and UCB datasets, including 377 concordantly upregulated and 2,245 concordantly downregulated genes, while 307 shared DEGs showed discordant directionality (Fig. 2E; Table S4). Thus, C/G-transformed UCB-HSPCs captured a directional patient-conserved C/G transcriptional core while retaining developmental and model-specific features.

To visualize whether concordantly upregulated genes occupied the leading edge of C/G-associated transcriptional states, we performed preranked GSEA using this shared upregulated gene set. The gene set was strongly enriched in both C/G⁺ pAML patient-ranked transcriptomes (NES = 7.19, P = 1.42 × 10⁻¹ ³) and C/G-transformed UCB-ranked transcriptomes (NES = 21.30, P = 1.27 × 10⁻¹ ³), indicating concentration among the most C/G-associated transcripts in both contexts (Fig. 2F,G; Table S5A). MSigDB Hallmark analysis identified enrichment of p53 pathway, IL2–STAT5 signaling, KRAS signaling, inflammatory response, and apoptosis programs (Fig. 2H; Table S5B). Together, these data support the C/G-transformed UCB-HSPC system as a developmentally relevant discovery model for early DNAm and chromatin analysis.

### C/G-transformed UCB-HSPCs show early methylation-linked active regulatory remodeling

Having established that C/G-transformed UCB-HSPCs capture an early patient-conserved transcriptional program, we asked whether this developmental system could reveal early DNAm and chromatin changes induced by the fusion. meEM-seq profiling showed broad DNAm differences between C/G-transformed UCB-HSPCs and GFP controls, comprising 4,662 hypermethylated and 24,093 hypomethylated DMRs. Promoters comprised 41.2% of hyper-DMRs, followed by intronic and distal intergenic regions. Hypomethylated DMRs were more broadly distributed across intronic, promoter, and distal intergenic compartments (Fig. 3A; Table S6A). We integrated UCB DMRs with patient DNAm and transcriptomic datasets to classify DMR-linked DEGs by cross-context detection. Most were UCB-specific (3,449/4,151; 83.1%), whereas 342 were patient-specific (8.2%) and 360 were detected in both contexts (8.7%; Fig. 3B; Table S6B). We term the latter “shared early-maintained” descriptively; these platform-dependent, cross-sectional categories do not establish absence from the other context or persistence within the same clone.

**Figure 3.**
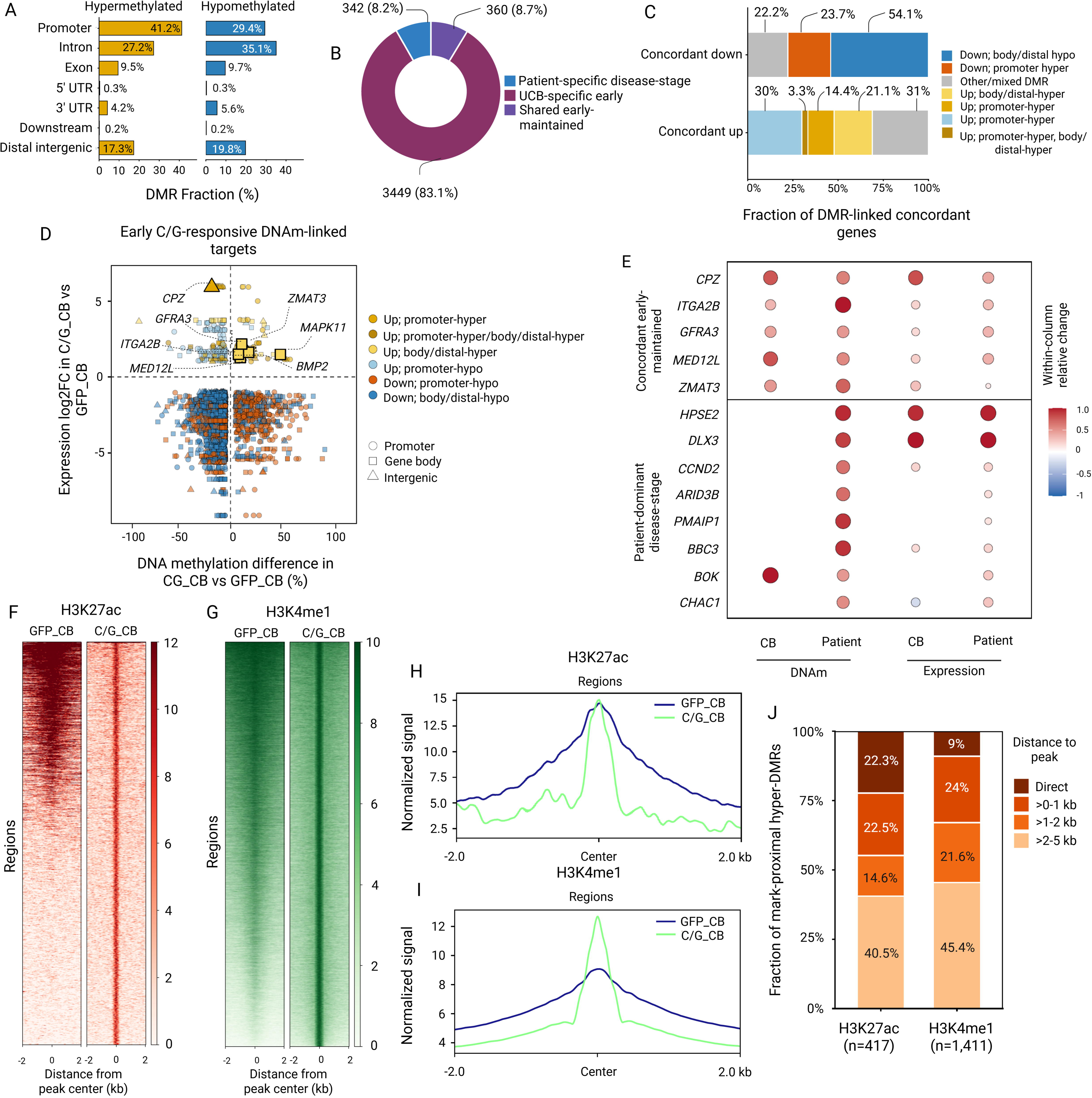
C/G-induced DNA methylation remodeling defines early DNAm-linked transcriptional programs that intersect active/primed chromatin. **(A)** Genomic distribution of C/G-induced 200-bp DMRs in C/G-transformed UCB-HSPCs versus GFP controls, stratified by hypermethylation or hypomethylation in C/G-transformed cells. Fractions were calculated within each DMR class. **(B)** Classification of DMR-linked DEGs as UCB-specific early, patient-specific disease-stage, or shared early-maintained, based on whether associated DNAm changes were detected in the C/G-transformed UCB system, patient leukemia, or both contexts. **(C)** Region-aware DNAm-expression patterns among DMR-linked concordantly upregulated or downregulated genes. Fractions were calculated within each concordant expression class. **(D)** DMR-level integration of DNAm and expression in C/G-transformed UCB-HSPCs. Each point represents a DMR linked to a concordantly regulated gene. Color indicates DNAm-expression class, and shape indicates genomic region. Selected early C/G-responsive DNAm-linked genes are labeled. **(E)** Dot-matrix summary of selected concordant early-maintained and patient-dominant disease-stage DNAm-linked genes. Columns show UCB DNAm, patient DNAm, UCB expression, and patient expression, with values scaled within each column for visualization. **(F,G)** H3K27ac and H3K4me1 CUT&RUN heatmaps centered on peak regions in GFP-control and C/G-transformed UCB-HSPCs. **(H,I)** Aggregate H3K27ac and H3K4me1 signal profiles across peak-centered regions. **(J)** Proximity of C/G-induced hyper-DMRs to H3K27ac- and H3K4me1-marked chromatin. Hyper-DMRs within 5 kb of each histone mark were binned by distance from the nearest peak. Fractions were calculated among mark-proximal hyper-DMRs. CB, cord blood; C/G, CBFA2T3–GLIS2; DNAm, DNA methylation; DMR, differentially methylated region; GFP, green fluorescent protein; H3K4me1, histone H3 lysine 4 monomethylation; H3K27ac, histone H3 lysine 27 acetylation; HSPC, hematopoietic stem/progenitor cell; pAML, pediatric acute myeloid leukemia; UCB, umbilical cord blood.

Region-aware integration of DNAm and expression revealed both canonical and noncanonical methylation-expression relationships. Among concordantly upregulated DMR-linked genes, 30.0% were associated with promoter hypomethylation, 14.4% with promoter hypermethylation, 21.1% with body/distal hypermethylation, and 3.3% with combined promoter and body/distal hypermethylation. Among concordantly downregulated DMR-linked genes, body/distal hypomethylation was dominant (54.1%), followed by promoter hypermethylation (23.7%) and other or mixed DMR configurations (22.2%; Fig. 3C,D; Table S6C). Representative early-maintained genes included CPZ, *ITGA2B*, *GFRA3*, *MED12L*, and *ZMAT3*, whereas *HPSE2*, *DLX3*, *CCND2*, *ARID3B*, *PMAIP1*, *BBC3*, *BOK*, and *CHAC1* showed stronger patient-dominant or disease-stage DNAm-expression features (Fig. 3E; Table S6D). These results show that C/G-induced DNAm-expression coupling includes expected promoter-linked patterns and noncanonical hypermethylated regulatory states associated with increased expression.

To determine whether C/G-induced hyper-DMRs occur near active or primed chromatin, we profiled H3K27ac and H3K4me1 in C/G-transformed and GFP-control UCB-HSPCs. Heatmap and aggregate-profile analyses showed focal redistribution of both marks at regulatory regions in C/G-transformed cells (Fig. 3F–I). H3K27ac profiling identified 1,755 regions, including 621 newly activated, 363 amplified, and 423 decreased regions; H3K4me1 profiling identified 42,443 regions, including 4,552 newly activated, 12,806 amplified, and 2,477 decreased regions (Fig. 3F-I; Table S6E). Consistent with active/primed enhancer-associated methylation, 417 and 1,411 hyper-DMRs were located within 5 kb of H3K27ac and H3K4me1 peaks, respectively. Among mark-proximal hyper-DMRs, 22.3% directly overlapped H3K27ac peaks and 9.0% overlapped H3K4me1 peaks, with most remaining regions located 1-5 kb from the nearest peak (Fig. 3J; Table S6F). H3K4me3 and H3K27me3 CUT&RUN showed C/G-associated promoter-associated and repressive chromatin redistribution, but did not define the dominant DNAm-linked CRE signal; therefore, downstream integration focused on H3K4me1/H3K27ac-marked active/primed regions (Fig. S1). Together, these discovery profiles place a subset of C/G-associated hyper-DMRs near active/primed chromatin, supporting spatial coupling between early methylation differences and active/primed CRE neighborhoods.

### Locus-level regulatory architecture distinguishes early-maintained and disease-stage C/G-linked loci

To determine whether the DMR-linked DEG classes identified by genome-wide DNAm-expression-chromatin integration were reflected at individual loci, we examined representative genes across the C/G-transformed UCB system and leukemia-stage C/G⁺ context. *CPZ*, *ITGA2B*, and *MAPK11* represented concordant early-maintained loci. In C/G-transformed UCB-HSPCs, these loci showed local hypermethylated DMRs near H3K4me1- and/or H3K27ac-marked regulatory regions, consistent with early C/G-induced remodeling of active or primed CREs (Fig. 4A,D,G). In the leukemia-stage context, M07e cells showed H3K4me1/H3K27ac-marked chromatin within the same gene loci, with patient hypermethylated CpGs detected in locus-proximal regulatory regions relative to normal HSC controls (Fig. 4B,E,H). These regulatory patterns were accompanied by increased *CPZ*, *ITGA2B*, and *MAPK11* expression in both the UCB system and primary C/G⁺ pAML (Fig. 4C,F,I). Thus, a subset of C/G-linked genes shows enhancer-proximal hypermethylation near active/primed chromatin in the UCB model, with related locus architecture also present in leukemia-stage datasets.

**Figure 4.**
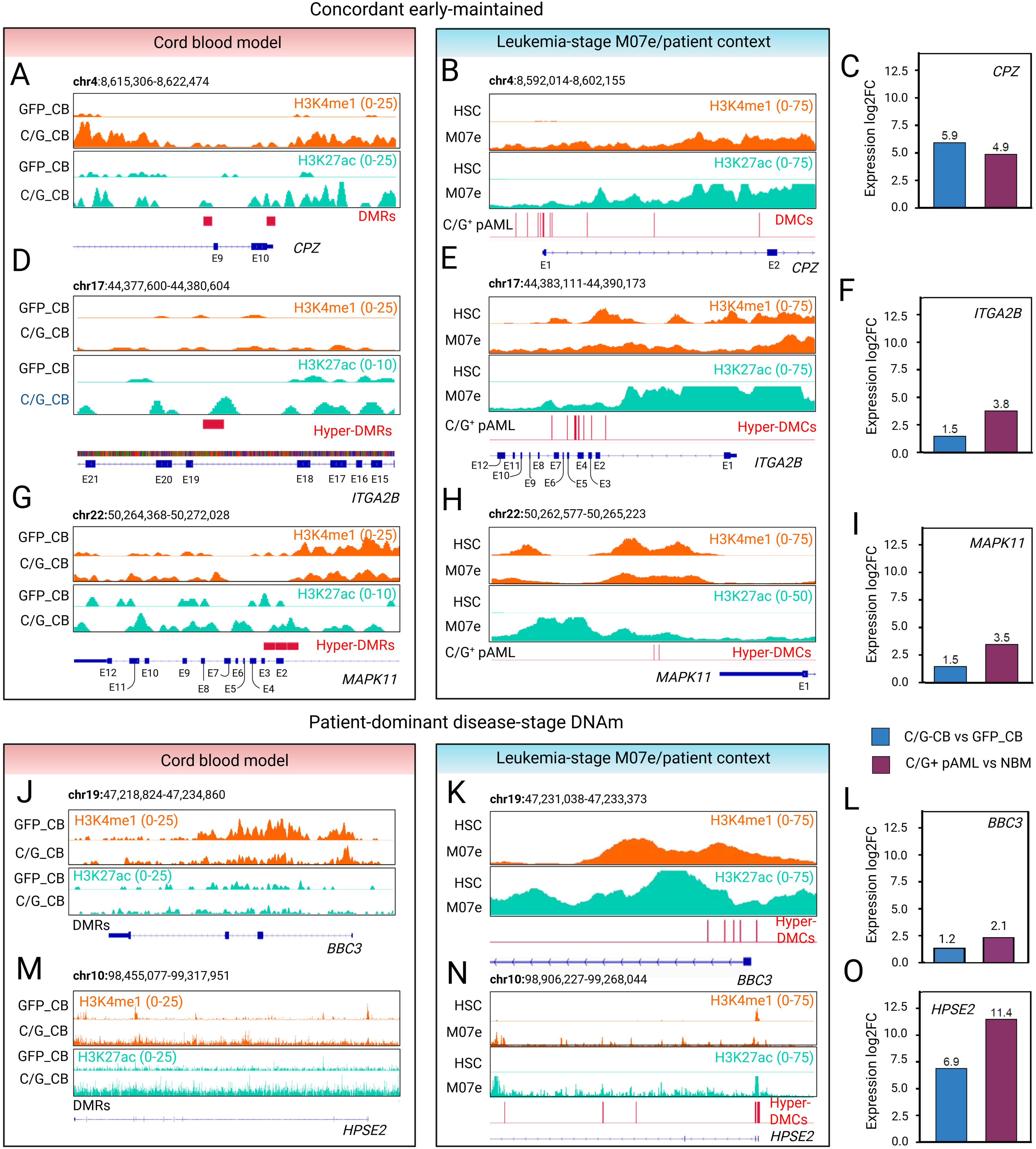
Locus-level regulatory architecture distinguishes early-maintained and disease-stage C/G-linked loci. **(A–C)** *CPZ* locus showing early-maintained enhancer-proximal hypermethylation, active/primed chromatin, and increased expression across the C/G-transformed UCB system and leukemia-stage C/G⁺ context. **(D–F)** *ITGA2B* locus showing early-maintained enhancer-proximal hypermethylation and active/primed chromatin remodeling across UCB and leukemia-stage contexts. **(G–I)** *MAPK11* locus showing early-maintained C/G-linked regulatory remodeling and increased expression across contexts. **(J–L)** *BBC3* locus showing patient-dominant disease-stage DNAm architecture, with limited C/G-induced hyper-DMR signal in the displayed UCB window but patient hypermethylated CpGs near H3K4me1/H3K27ac-marked regulatory regions in the leukemia-stage context. **(M–O)** *HPSE2* locus showing patient-dominant disease-stage DNAm architecture, with limited C/G-induced hyper-DMR signal in the displayed UCB window but prominent patient hypermethylated CpGs near active/primed chromatin and strong expression induction. For UCB panels, tracks compare GFP-transduced UCB-HSPCs with C/G-transformed UCB-HSPCs. For leukemia-stage panels, H3K4me1 and H3K27ac tracks compare normal HSC controls with M07e cells, with patient-derived C/G⁺ pAML hyper-DMCs overlaid. Red bars or vertical marks indicate C/G-induced hyper-DMRs in the UCB system or patient hyper-DMCs/DMR-like CpG clusters in the leukemia-stage context. Expression bars show log2FC for C/G-CB versus GFP-CB and C/G⁺ pAML versus NBM. Genome-browser tracks are shown in hg38 coordinates, with track ranges indicated in each panel. BBC3, BCL2-binding component 3; C/G, CBFA2T3–GLIS2; DMC, differentially methylated CpG; DMR, differentially methylated region; DNAm, DNA methylation; GFP, green fluorescent protein; H3K4me1, histone H3 lysine 4 monomethylation; H3K27ac, histone H3 lysine 27 acetylation; HPSE2, heparanase 2; HSC, hematopoietic stem cell; NBM, normal bone marrow; pAML, pediatric acute myeloid leukemia; UCB, umbilical cord blood.

We next examined *BBC3* and *HPSE2* as genes with patient-dominant disease-stage DNAm architecture. In the displayed UCB windows, these loci did not show prominent C/G-induced enhancer-proximal hyper-DMRs, despite C/G-associated expression changes (Fig. 4J,M). In contrast, the leukemia-stage C/G⁺ context showed patient hypermethylated CpGs positioned near H3K4me1/H3K27ac-marked regulatory regions in M07e (Fig. 4K,N). *BBC3* showed a modest expression increase in the UCB system and stronger expression in primary C/G⁺ pAML, whereas *HPSE2* was strongly upregulated in both contexts, with the highest expression increase in patient leukemia (Fig. 4L,O). Together, these examples show early-associated and disease-stage-prominent methylation-linked active/primed architectures, each associated with increased expression.

### Active/primed hypermethylated CREs are embedded in ERG/ETS-like motif-defined regulatory neighborhoods

Having shown that C/G-linked hyper-DMRs can reside near active or primed chromatin, we next asked whether these methylated CRE-associated regions share a regulatory motif architecture. We focused on 1,583 C/G-CB hyper-DMRs within 5 kb of H3K27ac or H3K4me1 peaks. HOMER known-motif analysis nominated ETS-family, AP-1/bZIP, HOX/MEIS, and CTCF/ZNF motif classes. Among these, an ERG/ETS-like motif emerged as an expression-supported candidate context because ERG was upregulated in the C/G-CB model, whereas SPIB was reduced (Fig. 5A; Table S7A). We therefore interpreted the ETS signal as ERG/ETS-like motif architecture rather than direct evidence of genome-wide ERG occupancy.

**Figure 5.**
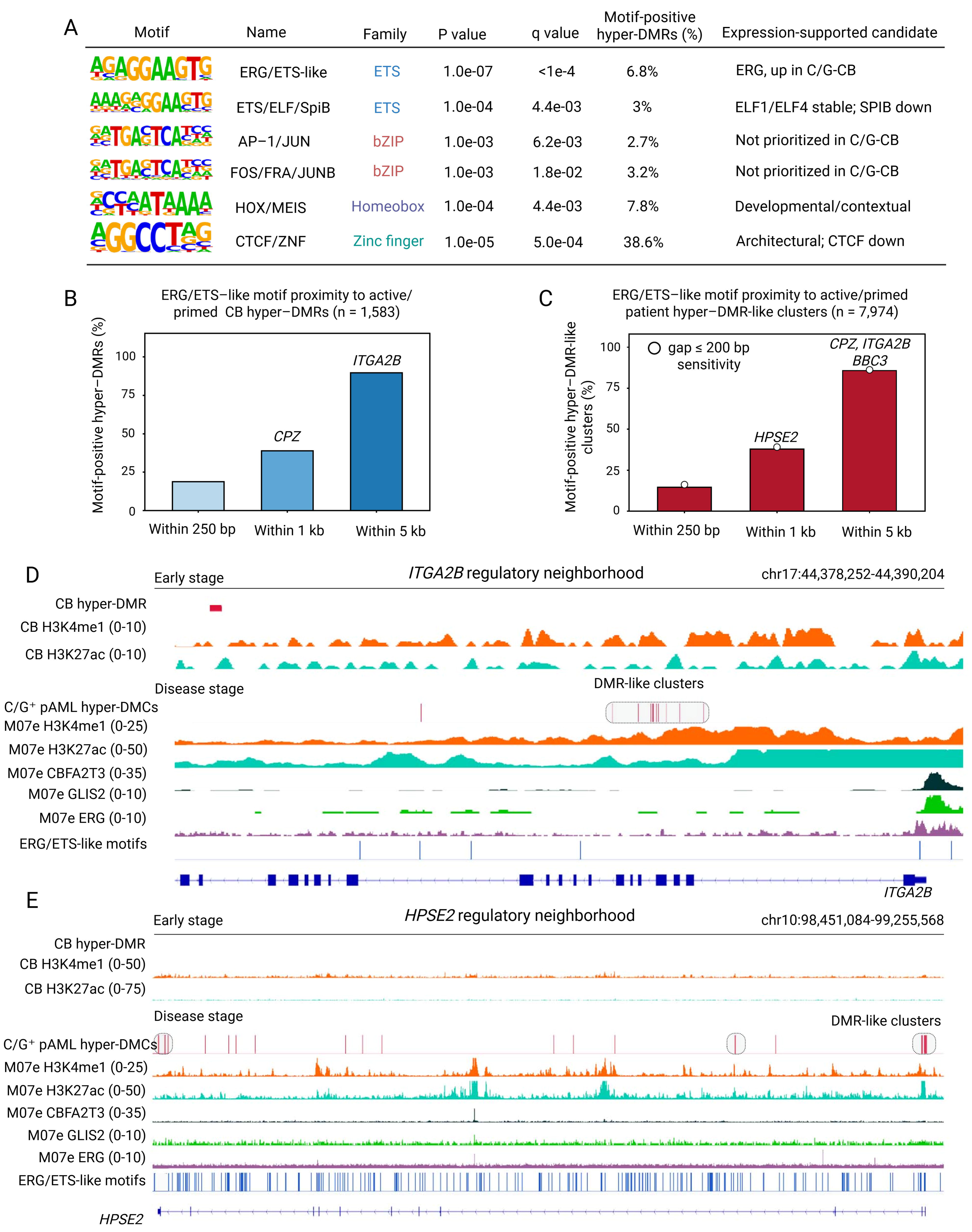
Active/primed hypermethylated CREs are embedded in ERG/ETS-like motif-defined regulatory neighborhoods. **(A)** Representative HOMER known-motif enrichment among active/primed C/G-transformed UCB hyper-DMRs, defined as C/G-induced hyper-DMRs located within 5 kb of H3K27ac or H3K4me1 peaks. ERG/ETS-like motifs were prioritized based on motif enrichment and expression-supported candidate interpretation. **(B)** ERG/ETS-like motif proximity to 1,583 active/primed C/G-transformed UCB hyper-DMRs. Bars show the fraction of hyper-DMRs with an ERG/ETS-like motif within 250 bp, 1 kb, or 5 kb. **(C)** ERG/ETS-like motif proximity to 7,974 active/primed patient hyper-DMR-like CpG clusters. Bars show motif proximity within 250 bp, 1 kb, or 5 kb. Open circles indicate the gap ≤200 bp cluster-definition sensitivity analysis. **(D)** *ITGA2B* regulatory neighborhood showing an early-stage C/G-CB hyper-DMR, active/primed chromatin, patient-derived leukemia-stage hyper-DMC/DMR-like clusters, M07e C/G occupancy, M07e ERG signal, and ERG/ETS-like motif sites. **(E)** *HPSE2* regulatory neighborhood showing limited early-stage hyper-DMR signal but prominent patient-derived leukemia-stage hyper-DMC/DMR-like clusters within an M07e active/primed chromatin environment containing ERG/ETS-like motif sites. For Panels D and E, early-stage tracks show C/G-transformed UCB hyper-DMRs and UCB H3K4me1/H3K27ac signal. Disease-stage tracks show M07e H3K4me1/H3K27ac, M07e CBFA2T3/GLIS2 occupancy, M07e ERG signal, and ERG/ETS-like motif sites, with patient-derived C/G⁺ pAML hyper-DMCs or DMR-like CpG clusters overlaid. C/G, CBFA2T3–GLIS2; CRE, cis-regulatory element; DMC, differentially methylated CpG; DMR, differentially methylated region; ERG, ETS-related gene; H3K27ac, histone H3 lysine 27 acetylation; H3K4me1, histone H3 lysine 4 monomethylation; HPSE2, heparanase 2; UCB, umbilical cord blood.

To determine whether ERG/ETS-like motifs were spatially positioned near active/primed hyper-DMRs, we scanned hg38 for the top ETS-like motif and calculated motif proximity to the 1,583 active/primed C/G-CB hyper-DMRs. ERG/ETS-like motifs were detected within 250 bp of 297 hyper-DMRs (18.8%), within 1 kb of 615 hyper-DMRs (38.9%), and within 5 kb of 1,420 hyper-DMRs (89.7%; Fig. 5B; Table S7B). We next performed the same analysis in the disease-stage context using patient hypermethylated DMCs collapsed into DMR-like CpG clusters and restricted to clusters within 5 kb of M07e H3K27ac or H3K4me1 peaks. Among 7,974 active/primed patient hyper-DMR-like clusters, ERG/ETS-like motifs were detected within 250 bp of 1,162 clusters (14.6%), within 1 kb of 3,021 clusters (37.9%), and within 5 kb of 6,835 clusters (85.7%; Fig. 5C; Table S7C). Similar results using a broader DMR-like clustering definition supported robustness to the regional-clustering threshold.

Locus-level visualization further supported this motif-defined architecture. At ITGA2B, an early-stage C/G-CB hyper-DMR was positioned within an H3K4me1/H3K27ac-marked regulatory neighborhood that also showed patient-derived leukemia-stage hyper-DMC clusters, M07e C/G occupancy, ERG signal, and nearby ERG/ETS-like motif sites (Fig. 5D). In contrast, HPSE2 showed limited early-stage hyper-DMR signal but prominent patient-derived leukemia-stage hyper-DMC clusters within an M07e active/primed chromatin environment containing ERG/ETS-like motif sites (Fig. 5E). Together, these analyses show recurrent spatial co-occurrence of C/G-associated active/primed hypermethylated regions with ERG/ETS-like motif architecture across early and leukemia-stage contexts, without establishing genome-wide ERG occupancy or dependence.

### DNMT3B supports a selected C/G-linked methylation-expression state without altering bulk 5mC

To test whether C/G promotes DNMT3B as part of its methylation-linked program, we expressed FLAG-tagged GFP, intact C/G, or a C/G mutant lacking NHR2 and ZF2 to ZF4 in HEK-293T cells (Fig. 6A). FLAG immunoblotting confirmed construct expression (Fig. 6B). Intact C/G most clearly increased DNMT3B protein abundance among the DNMT/TET/UHRF1 regulators examined, compared with GFP and mutant C/G controls (Fig. 6C), consistent with an association between intact C/G expression and DNMT3B induction in this heterologous comparison. In endogenous C/G⁺ M07e cells, DNMT3B KO depleted DNMT3B protein and gRNA-resistant rescue restored DNMT3B abundance (Fig. 6D). Global 5mC was not significantly altered across WT, DNMT3B-KO, and rescue cells (Fig. 6E), indicating that DNMT3B loss did not measurably affect bulk 5mC under these conditions and motivating locus-level analysis.

**Figure 6.**
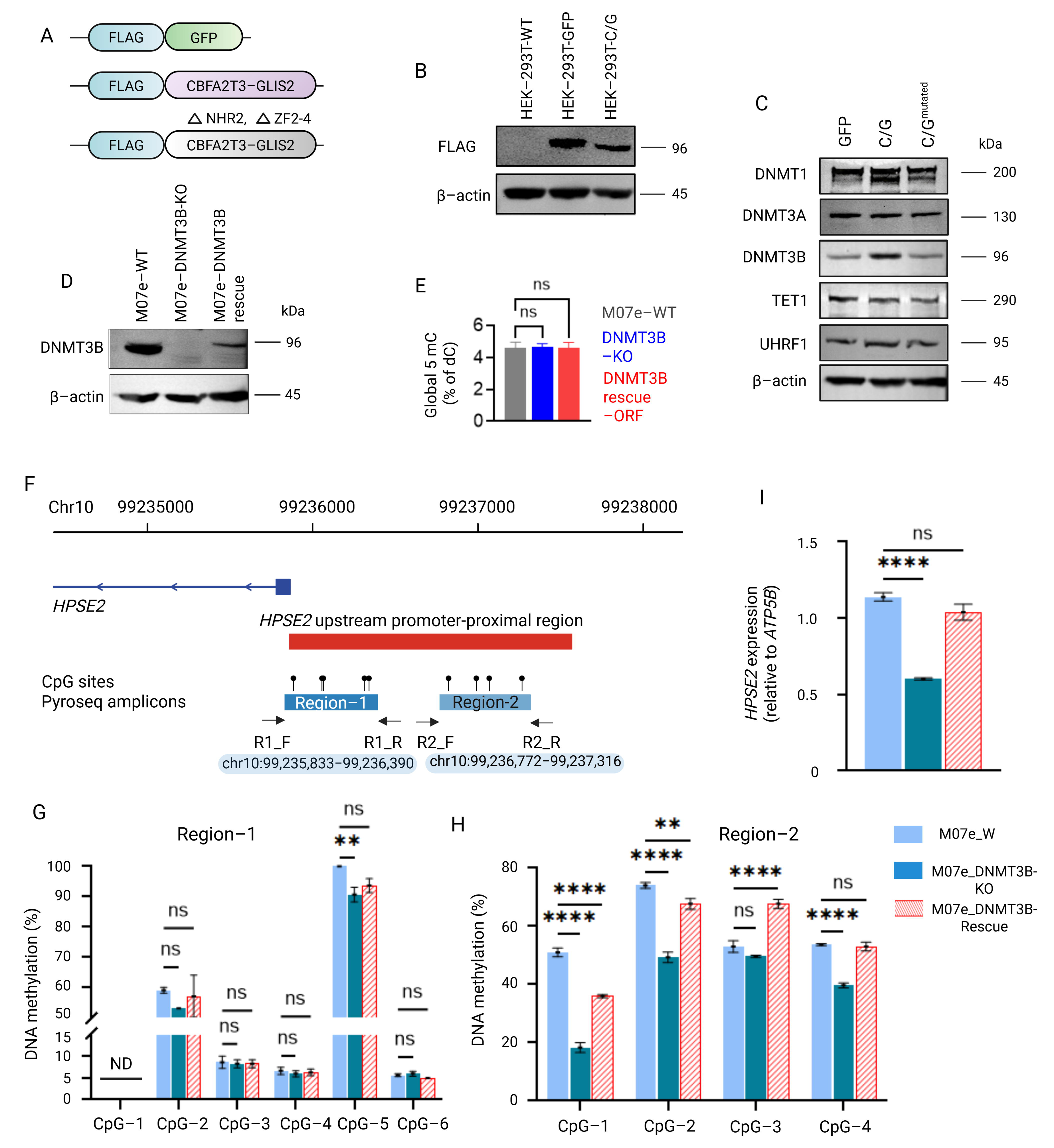
DNMT3B supports a selected C/G-linked methylation-expression state without altering bulk 5mC. **(A)** Schematic of FLAG-tagged constructs used to compare DNMT3B abundance after expression of GFP control, intact CBFA2T3–GLIS2 (C/G), or a C/G mutant lacking the CBFA2T3 NHR2 domain and GLIS2 ZF2 to ZF4 domains. **(B)** FLAG immunoblot analysis confirming expression of FLAG-tagged constructs in HEK-293T cells, with β-actin as loading control. **(C)** Immunoblot analysis of DNMT1, DNMT3A, DNMT3B, TET1, and UHRF1 in HEK-293T cells expressing GFP, intact C/G, or mutant C/G. β-actin was used as loading control. **(D)** Immunoblot validation of DNMT3B depletion in M07e DNMT3B-KO cells and gRNA-resistant DNMT3B rescue. β-actin was used as loading control. **(E)** Global 5mC measured by LC-MS in M07e WT, DNMT3B-KO, and DNMT3B-rescue cells. Values are shown as percent 5mC relative to deoxycytidine. **(F)** Schematic of HPSE2 upstream promoter-proximal regions assayed by pyrosequencing. CpG sites and amplicons for Region 1 and Region 2 are shown in hg38 coordinates. **(G,H)** CpG-level DNA methylation measured by pyrosequencing across HPSE2 Region 1 and Region 2 in M07e WT, DNMT3B-KO, and DNMT3B-rescue cells. **(I)** HPSE2 expression measured by RT-qPCR in M07e WT, DNMT3B-KO, and DNMT3B-rescue cells. C/G, CBFA2T3–GLIS2; DNAm, DNA methylation; DNMT, DNA methyltransferase; HEK-293T, human embryonic kidney 293T; KO, knockout; LC-MS, liquid chromatography-mass spectrometry; ns, not significant; ORF, open reading frame; TET, ten-eleven translocation methylcytosine dioxygenase; UHRF1, ubiquitin-like with PHD and ring finger domains 1; WT, wild type. Figure 6, panel A was created with BioRender.com under publication license SO29XM8W78 (Created in BioRender. Roy choudhury, S. (2026) https://BioRender.com/90tv02x)

Given prior genome-wide evidence that *DNMT3B* loss alters multiple DNAm-linked transcriptional programs [10, 11], we selected *HPSE2* for rescue-based validation because it showed strong C/G-associated expression and patient-stage promoter-proximal hypermethylation. Pyrosequencing across two upstream regions showed CpG-selective methylation loss after *DNMT3B* KO, with restoration toward WT after rescue (Fig. 6F-H). *HPSE2* expression similarly decreased after DNMT3B KO and was restored by rescue (Fig. 6I). Together, these findings provide locus-level evidence that *DNMT3B* contributes to a selected methylation-expression state despite stable bulk 5mC.

### DNMT3B perturbation lowers the venetoclax response threshold in C/G⁺ AML

Because BH3-mimetic activity has been reported in C/G leukemia but VEN response was incomplete in our models, we tested whether DNMT3B status alters the VEN response threshold [12, 13]. Genetic DNMT3B loss shifted the M07e VEN IC50 from approximately 1000 to 327 nM (Fig. 7A). Treatment with the DNMT3B-biased tool compound Nan-A similarly shifted the apparent VEN IC50 from approximately 1000 to 130 nM in M07e cells and from approximately 500 to 65 nM in WSU-AML cells (Fig. 7B).

**Figure 7.**
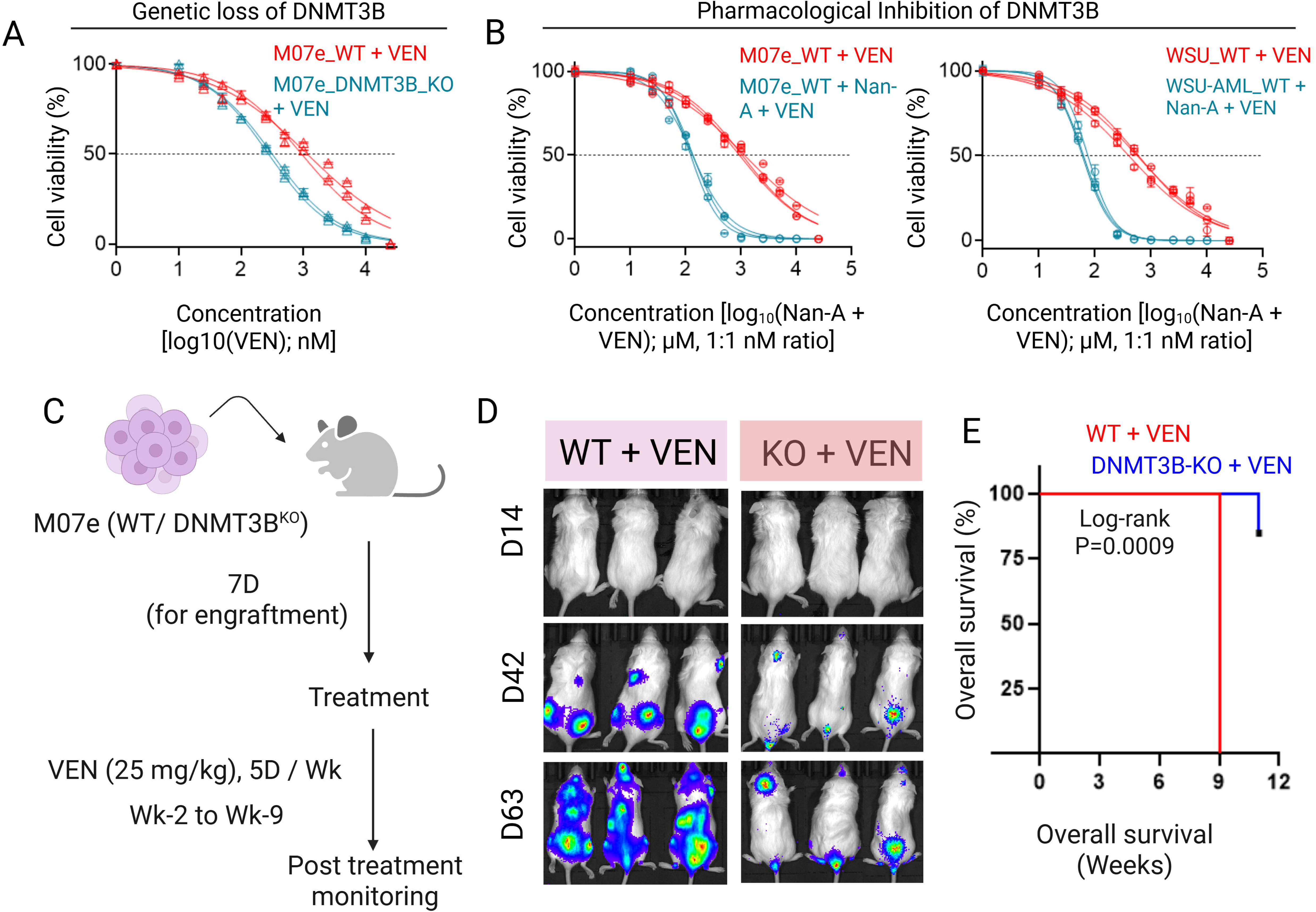
DNMT3B perturbation lowers the venetoclax response threshold in C/G⁺ AML. **(A)** Cell-viability response to venetoclax in M07e WT and DNMT3B-KO cells. DNMT3B loss shifted the venetoclax response curve relative to WT cells. **(B)** Cell-viability response to venetoclax alone or Nanaomycin-A plus venetoclax in C/G⁺ AML cell lines. M07e and WSU-AML cells were treated with venetoclax alone or combined with Nanaomycin-A at a 1:1 ratio for 48 h. **(C)** In vivo treatment schema for M07e WT and DNMT3B-KO xenografts treated with venetoclax. NSG-SGM3 mice were injected by tail vein, allowed to engraft for 7 days, and treated with venetoclax at 25 mg/kg, 5 days per week, from week 2 to week 9. **(D)** Serial bioluminescence imaging of male and female mice bearing M07e WT or DNMT3B-KO xenografts during venetoclax treatment. **(E)** Kaplan-Meier survival analysis of venetoclax-treated M07e WT and DNMT3B-KO xenografts. Statistical significance was assessed by log-rank test. All xenografts received VEN; this comparison therefore evaluates genotype-associated disease control under VEN exposure rather than a formal genotype-by-treatment interaction. C/G, CBFA2T3–GLIS2; DNMT, DNA methyltransferase; KO, knockout; Nan-A, Nanaomycin-A; NSG-SGM3, NOD.Cg-Prkdcscid Il2rgtm1Wjl Tg(CMV-IL3,CSF2,KITLG)1Eav/MloySzJ; VEN, venetoclax; WT, wild type. Figure 7, panel C was created with BioRender.com under publication license ZS29XM9J3H (Created in BioRender. Roy choudhury, S. (2026) https://BioRender.com/vh966iv)

We compared VEN-treated M07e WT and DNMT3B-KO xenografts (Fig. 7C). DNMT3B-KO xenografts showed lower BLI signal during treatment and prolonged survival relative to WT xenografts (log-rank P = 0.0009; Fig. 7D). Because untreated genotype controls were not included, this experiment demonstrates improved disease control under VEN exposure but does not resolve a genotype-by-treatment interaction.

### HMAs enhance venetoclax response and are associated with apoptosis-related transcriptional and mitochondrial changes in C/G⁺ AML

Because DNMT3B loss did not alter bulk 5mC, we asked whether maintenance machinery was increased after DNMT3B perturbation. DNMT3B KO increased DNMT1 and UHRF1 abundance relative to WT and scramble controls, whereas DNMT3A remained largely unchanged (Fig. 8A). DNMT1 was recovered in UHRF1 immunoprecipitates from DNMT3B-KO cells, consistent with DNMT1-UHRF1 association after DNMT3B loss (Fig. 8B). These observations prompted us to test clinically used HMAs, which perturb DNMT-dependent maintenance more broadly than DNMT3B loss, as a strategy to lower the VEN response threshold (Fig. 8C).

**Figure 8.**
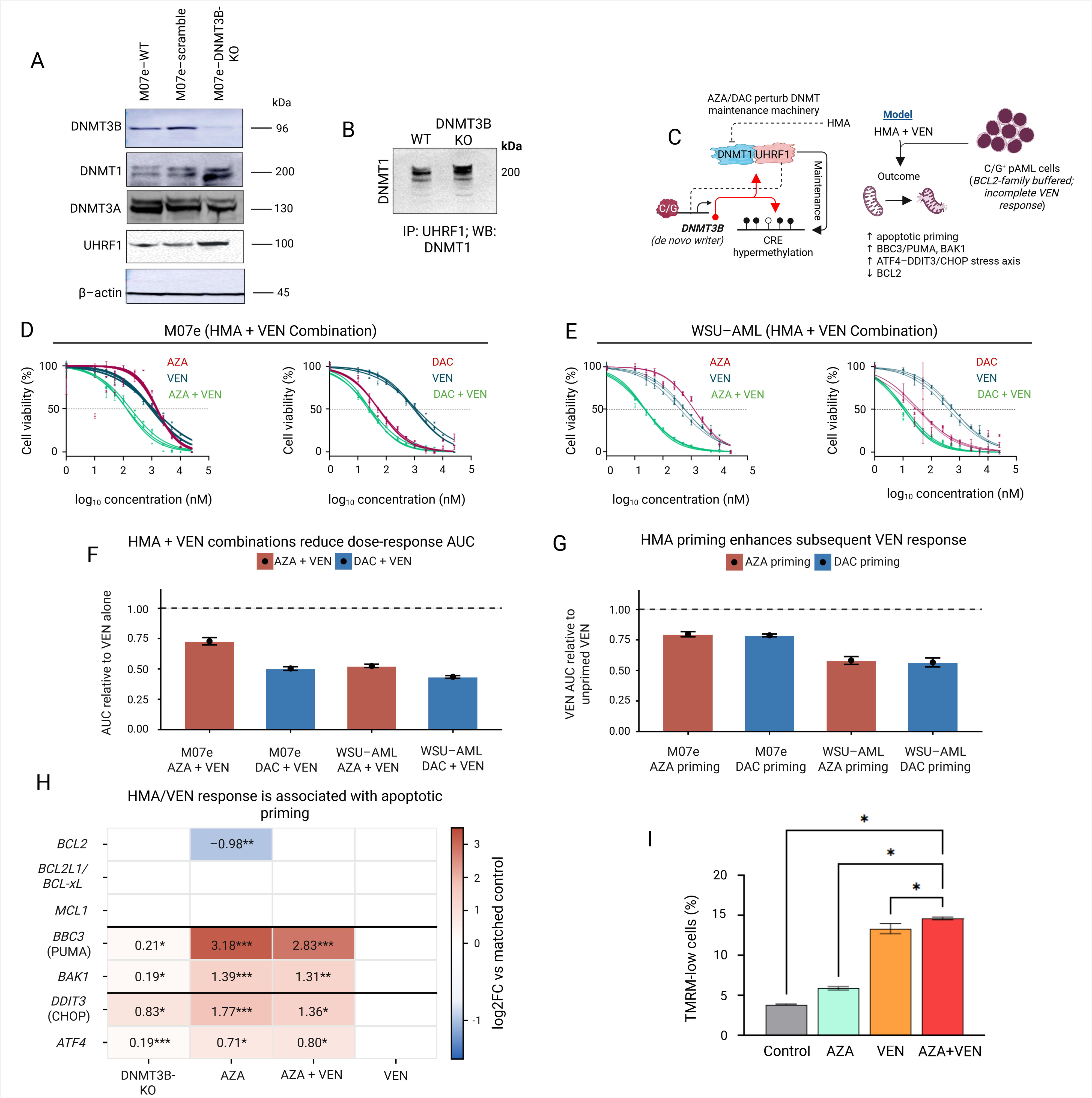
HMAs enhance venetoclax response and are associated with apoptotic priming in C/G⁺ AML. **(A)** Immunoblot analysis of DNMT3B, DNMT1, DNMT3A, and UHRF1 in M07e WT, scramble-control, and DNMT3B-KO cells. β-actin was used as loading control. **(B)** DNMT1 detection after UHRF1 co-immunoprecipitation in M07e WT and DNMT3B-KO cells. **(C)** Working model in which DNMT3B loss is accompanied by increased DNMT1/UHRF1 abundance and DNMT1-UHRF1 association, whereas HMAs perturb DNMT-dependent maintenance more broadly and lower the VEN response threshold. **(D,E)** Cell-viability response to AZA, DAC, VEN, or equimolar HMA+VEN combinations in M07e and WSU-AML cells. For combination curves, HMA and VEN were combined at equimolar concentrations. **(F)** Dose-response AUC for concurrent HMA+VEN treatment normalized to VEN alone in M07e and WSU-AML cells. The dashed line indicates the VEN-alone reference. **(G)** VEN dose-response AUC after AZA or DAC priming, normalized to unprimed VEN response in M07e and WSU-AML cells. The dashed line indicates the unprimed VEN reference. **(H)** Focused apoptosis-related RNA-seq heatmap in M07e cells after DNMT3B KO, AZA, AZA+VEN, or VEN treatment. Values show log2 fold-change versus matched control. Asterisks indicate statistically significant differential expression, and blank cells indicate non-significant changes in this focused panel. **(I)** TMRM flow cytometry showing the fraction of TMRM-low M07e cells after control, AZA, VEN, or AZA+VEN treatment. Increased TMRM-low fraction indicates mitochondrial membrane depolarization. AZA, azacitidine; C/G, CBFA2T3–GLIS2; DAC, decitabine; DNMT, DNA methyltransferase; HMA, hypomethylating agent; IP, immunoprecipitation; KO, knockout; TMRM, tetramethylrhodamine methyl ester; UHRF1, ubiquitin-like with PHD and ring finger domains 1; VEN, venetoclax; WB, immunoblot; WT, wild type. Figure 8, panel C was created with BioRender.com under publication license ZS29XM9J3H Created in BioRender. Roy choudhury, S. (2026) https://BioRender.com/6osru1m)

We then tested whether HMAs enhance VEN response in endogenous C/G⁺ AML models. In M07e cells, AZA, DAC, and VEN produced dose-dependent viability reductions with approximate IC50 values of 1500, 50, and 1000 nM, respectively. Equimolar AZA+VEN and DAC+VEN further reduced the apparent IC50 to approximately 150 and 25 nM, respectively, calculated at the concentration of each agent in the combination (Fig. 8D). In WSU-AML cells, AZA, DAC, and VEN IC50 values were approximately 1250, 30, and 500 nM, respectively; equimolar AZA+VEN and DAC+VEN reduced the apparent IC50 to approximately 25 and 12 nM (Fig. 8E). Concurrent HMA+VEN reduced dose-response AUC relative to VEN alone in M07e cells (AZA+VEN, 0.73 ± 0.03; DAC+VEN, 0.50 ± 0.02) and WSU-AML cells (AZA+VEN, 0.52 ± 0.01; DAC+VEN, 0.43 ± 0.01; Fig. 8F). HMA priming also enhanced subsequent VEN response, reducing normalized VEN AUC to 0.80 ± 0.02 and 0.79 ± 0.01 after AZA or DAC priming in M07e cells and to 0.58 ± 0.03 and 0.57 ± 0.04 in WSU-AML cells, respectively (Fig. 8G).

Because AZA perturbs DNMT-dependent methylation maintenance more broadly than DNMT3B loss, we asked whether genetic DNMT3B perturbation and AZA-based treatment converge on apoptosis-related transcriptional responses. Focused RNA-seq analysis showed selected apoptotic/stress-response induction after DNMT3B KO, broader induction of *BBC3*/PUMA, *BAK1*, *DDIT3*/CHOP, and *ATF4* after AZA or AZA+VEN, and limited transcriptional priming after VEN alone (Fig. 8H). TMRM flow cytometry showed increased mitochondrial depolarization with VEN-containing treatment, with the highest TMRM-low fraction after AZA+VEN (Fig. 8I). Together, these data associate broader DNMT perturbation with a VEN-responsive state characterized by apoptotic/stress transcription and mitochondrial depolarization.

## Discussion

This study identifies transcriptionally permissive enhancer-proximal hypermethylation as a recurrent regulatory feature of C/G-positive pAML. C/G leukemia is established as an enhancer-rewiring disease in which fusion occupancy creates aberrant CRE activity and leukemic transcriptional identity [8, 9]. Our data extend this framework by showing that C/G-associated methylation remodeling can coexist with H3K27ac/H3K4me1-marked chromatin and increased expression [10]. Importantly, only a focused fraction of patient hypermethylated CpGs overlapped M07e C/G occupancy, arguing that direct fusion binding is not the sole explanation and that secondary regulatory mechanisms likely propagate the broader DNAm landscape. Thus, DNAm is best viewed here as a candidate regulatory layer coupled to, rather than synonymous with, fusion occupancy [10, 14].

The UCB-HSPC model provides developmental timing but does not reproduce the full cellular heterogeneity or leukemia-stage chromatin of primary C/G-positive AML. We therefore used it to nominate early fusion-associated events and patient methylation, leukemia-stage chromatin, and expression data to identify cross-context support [5, 6]. The predominance of UCB-specific DMR-linked genes indicates extensive model-or stage-restricted remodeling, whereas the smaller shared class marks features detectable in both early and leukemia-stage datasets. Because these comparisons are cross-sectional, “early-maintained” denotes detection in both contexts rather than longitudinal persistence within the same leukemic clone. Patient DNAm, M07e chromatin/occupancy, and UCB maps also derive from different specimens, so their integration establishes cross-context concordance rather than same-cell co-occupancy [15].

Our data support DNMT3B as one contributor to selected methylation-linked regulatory states, not as the sole methyltransferase governing the C/G epigenome. Intact C/G most prominently increased DNMT3B among the regulators examined in a heterologous expression system, while prior occupancy data place C/G at the DNMT3B promoter [10, 16-18]. These observations are consistent with, but do not by themselves prove, direct promoter activation. DNMT3B loss left bulk 5mC unchanged, arguing against a global methylation-collapse mechanism. Instead, HPSE2 rescue provides locus-level evidence that DNMT3B can support CpG methylation and expression at a selected C/G-associated locus, consistent with prior genome-wide perturbation studies [11, 19]. DNMT3A, DNMT1, TET enzymes, and other C/G-regulated factors may contribute to the remaining regional and global methylation changes.

Increased DNMT1 and UHRF1 abundance after DNMT3B loss, together with recovery of DNMT1 in UHRF1 immunoprecipitates, supports the possibility that maintenance machinery contributes to preservation of bulk 5mC. These data do not establish functional compensation or identical genomic targeting by DNMT1 and DNMT3B. Rather, they motivate a model in which replication-coupled maintenance can preserve overall methylation while selected DNMT3B-linked loci remain altered [20, 21].

ERG/ETS-like motifs were frequently proximal to active/primed hyper-DMRs in both UCB and patient-stage datasets. This convergence identifies ETS-like sequence architecture as a recurrent context in which methylation and active chromatin co-occur [8]. Because ETS motifs are common and motif proximity is not occupancy, these analyses do not establish genome-wide ERG binding, preferential enrichment relative to all active enhancers, or functional ERG dependence. The locus tracks provide contextual support, but enhancer or ERG-family perturbation would be required to test causality.

Functionally, DNMT3B loss lowered the VEN response threshold in vitro, and DNMT3B-deficient xenografts showed improved disease control during VEN treatment [22, 23]. Because untreated genotype controls were not included, the xenograft experiment cannot distinguish reduced baseline leukemic fitness from a genotype-by-VEN interaction. Prior reports support BH3-mimetic activity and HMA+VEN feasibility in C/G AML [12, 13]. Nan-A provides supportive pharmacologic evidence but should be interpreted as a DNMT3B-biased tool compound rather than a selective inhibitor. AZA and DAC act more broadly and therefore do not phenocopy DNMT3B loss. The translational contribution is not the established HMA+VEN combination itself, but identification of a C/G-associated DNMT-supported state linked to VEN response. The accompanying apoptotic/stress transcription and TMRM loss are consistent with apoptosis-related pathway activation and mitochondrial depolarization as correlates of response, not a fully resolved causal mechanism [24, 25].

## Conclusion

Together, these findings define C/G-positive AML as an enhancer-rewired leukemia with transcriptionally permissive hypermethylated CRE states. These states are detected in the early UCB model, involve DNMT3B at selected loci, and are linked to a lower venetoclax response threshold after DNMT perturbation, providing a biological rationale for evaluating HMA-based combinations in this high-risk subtype.

## Data availability

Publicly available datasets used in this study include TARGET/COG pediatric AML RNA-seq data from dbGaP accession phs000465.v19.p8, patient DNA methylation datasets from GEO accessions GSE124413 and GSE190931, and RNA-seq data from C/G-transformed and GFP-transduced UCB-CD34⁺ cells from GEO accession GSE181726. Previously generated C/G occupancy and leukemia-stage chromatin datasets were analyzed as described in the Methods and Supplementary Information. Newly generated CUT&RUN (GSE338527) and meEM-seq (GSE338586) datasets from the C/G-transformed cord blood model have been deposited in GEO. Source data supporting the figures are provided with the manuscript and Supplementary Tables. Custom plasmid constructs generated in this study are available from the corresponding author upon reasonable request, subject to institutional material transfer agreement and any applicable restrictions. Additional processed data are available from the corresponding author upon reasonable request.

## Acknowledgements

We thank Ms. Andrea Harris at the UAMS Flow Cytometry Core for technical assistance with flow cytometry acquisition and analysis. Schematic illustrations were created with BioRender.com.

## Contributions

SRC conceived the study and designed the experiments. AC, RR and PB performed the experimental work. RR and SM generated the UCB-GFP and UCB-C/G cord blood models. AC and SRC analyzed the data, with input from JF. PB performed the in vivo DNMT3B experiment and UHRF1 immunoprecipitation studies. SRC supervised the project and drafted the manuscript. All authors critically reviewed and approved the final version of the manuscript.

## Ethics declaration Competing interests

The authors declare no competing interests.

## Ethics approval

DNA methylation array analyses were performed using specimens from consented patients enrolled in the TARGET-AML study (phs000465). These controlled-access genomic datasets are available through the database of Genotypes and Phenotypes (dbGaP) and the NCI Genomic Data Commons (GDC). The present study used de-identified secondary genomic data and did not involve prospective enrollment of human subjects. Cord blood CD34⁺ cells used to generate the UCB-GFP and UCB-C/G models were obtained through the Fred Hutch Cancer Biobank/Fred Hutchinson Cancer Center under institutional approvals and written informed consent from donors, as previously described. The animal studies described in this manuscript were approved by the Institutional Animal Care and Use Committee (IACUC), IACUC ID: IPROTO202200000037.

