## Supplementary Information for "CBFA2T3–GLIS2 establishes transcriptionally permissive enhancer hypermethylation associated with venetoclax response in pediatric AML"

### Supplementary Methods

*Patient cohorts, subtype annotation, and survival analysis*

Pediatric AML subtype distribution and survival analyses were performed using the TARGET/COG pAML cohort and associated clinical and molecular annotations. Cases were classified by French–American–British (FAB) subtype and by major molecular/fusion-defined group, including CBFA2T3–GLIS2, CBFB–MYH11, RUNX1–RUNX1T1, KMT2A-r AML, and AML lacking the indicated major fusions or classified as other AML. Survival groups shown in Fig. 1 included CBFA2T3–GLIS2 (n = 40), CBFB–MYH11 (n = 106), RUNX1–RUNX1T1 (n = 148), KMT2A-r AML (n = 196), and AML without the indicated major fusions ("None"; n = 326). CBFA2T3–GLIS2+ cases were further annotated by FAB subtype to assess representation across M7/acute megakaryoblastic leukemia and non-M7 AML. Overall survival was calculated from diagnosis to death or last follow-up and compared by Kaplan–Meier analysis with log-rank testing.

For patient transcriptomic analyses, C/G+ pAML and normal bone marrow (NBM) RNA-seq data were obtained from published RNA-seq datasets available through dbGaP (phs000465.v19.p8) [1]. Patient and NBM DNAm data were obtained from GEO accessions GSE124413 and GSE190931. Dataset accessions, sample inclusion criteria, normalization, differential expression, and methylation analyses are described in the RNA-seq and DNAm sections below.

*Cell lines and culture conditions*

C/G-positive AML cell lines M07e and WSU-AML were used as endogenous C/G+ leukemia models. C/G-negative AML cell lines MV4-11 and AML-193 were used as fusion-negative controls where indicated. M07e cells (ACC 104, DSMZ) were maintained in RPMI 1640 supplemented with 20% heat-inactivated FBS and recombinant human GM-CSF (10 ng/ml). WSU-AML cells (BioIVT) were maintained in RPMI 1640 supplemented with 20% heat-inactivated FBS. MV4-11 cells (CRL-9591, ATCC) were maintained in

RPMI 1640 supplemented with 10–20% heat-inactivated FBS. AML-193 cells (CRL-9589, ATCC) were maintained in 80% Iscove's modified Dulbecco's medium and 20% heat-inactivated FBS supplemented with recombinant human GM-CSF (10 ng/ml) and IL-3 (10 ng/ml). Cells were cultured at 37 °C with 5% $CO_2$ and maintained in logarithmic growth before molecular, immunoblotting, or drug-response assays. Cell lines were authenticated by short tandem repeat (STR) profiling and tested negative for mycoplasma before use. Cells were used within five passages after thawing.

*Immunoblotting and DNMT1-UHRF1 co-immunoprecipitation*

Whole-cell lysates were prepared in RIPA buffer containing protease and phosphatase inhibitors. Protein lysates were resolved by SDS-PAGE using NuPAGE 12% Bis-Tris Mini Protein Gels (#NP0342BOX, Thermo Scientific) and transferred to 0.45-µm PVDF membranes (#88518, Thermo Scientific). The CBFA2T3-GLIS2 fusion protein was detected using an antibody recognizing the N-terminal CBFA2T3 portion of the fusion (#ARP38578_T100, Aviva). DNMT1, DNMT3A, DNMT3B, UHRF1, TET-family proteins where indicated, FLAG, and β-actin were analyzed by standard immunoblotting using the antibodies listed in Table S8B. For whole-cell immunoblotting, UHRF1 was detected using anti-UHRF1 antibody (#12387, Cell Signaling Technology). β-actin was used as loading control. Immunoblot images were cropped for figure presentation, and molecular weight markers are indicated in kDa. For DNMT1-UHRF1 co-immunoprecipitation, lysates from M07e WT and DNMT3B-KO cells were prepared and processed using the Dynabeads Co-Immunoprecipitation Kit (#14321D, Thermo Fisher Scientific) according to the manufacturer's protocol. UHRF1-containing complexes were immunoprecipitated using recombinant anti-UHRF1 antibody [EPR18803-11] (#ab213223, Abcam). Co-precipitated DNMT1 was detected by immunoblotting using anti-DNMT1 antibody (#5032, Cell Signaling Technology). Input and immunoprecipitated fractions were resolved by SDS-PAGE and analyzed by immunoblotting as described above.

*C/G-transformed cord blood HSPC model*

C/G-transformed and GFP-transduced umbilical cord blood-derived CD34-positive hematopoietic stem and progenitor cells were generated using previously described lentiviral and endothelial co-culture methods [1, 2]. Briefly, UCB-derived CD34-positive HSPCs were transduced with lentiviral vectors expressing CBFA2T3–GLIS2/GFP or GFP control. Transduced GFP-positive cells were sorted and maintained on endothelial co-culture to support expansion and transformation-associated phenotypes. C/G-transformed and GFP-control cultures were harvested at day 42 for downstream flow cytometry, RNA-seq, DNAm profiling, and chromatin profiling. For the C/G-transformed cord blood model, RNA-seq data from C/G–GFP- and GFP-transduced UCB-CD34+ cells were obtained from GSE181726. At day 42, GFP-control and C/G-transformed cord blood cultures were analyzed by flow cytometry for CD34 (#130-120-519, Miltenyi Biotec) and FOLR1 (#130-129-539, Miltenyi Biotech) expression. FOLR1 was used as a C/G-restricted pAML-associated surface marker.

*EPIC methylation array processing and DMC annotation*

Patient/NBM DNAm data were analyzed from EPIC methylation array profiles comparing C/G+ pediatric AML samples with normal bone marrow controls. CpGs passing quality control were used as the tested background universe. Differentially methylated CpGs were defined using an absolute methylation difference threshold of at least 5% between C/G+ pAML and normal bone marrow, unless otherwise specified. CpGs with positive delta beta values were classified as hypermethylated in C/G+ pAML, whereas CpGs with negative delta beta values were classified as hypomethylated. Probe coordinates were converted to BED format using hg38 genomic coordinates, with CpG positions represented as 1-bp intervals. CpGs were annotated as promoter, gene body, or intergenic based on the annotation provided in the EPIC methylation analysis. Promoter-associated CpGs were defined according to the EPIC annotation and, where needed, cross-checked against transcription start site-proximal annotations.

*Histone CUT&RUN in C/G-transformed cord blood cells*

Day 42 C/G-transformed cord blood cultures were FACS-sorted for the FOLR1$^+$ C/G-associated fraction before chromatin profiling. GFP-transduced cord blood controls were processed in parallel. Histone CUT&RUN was performed using $1 \times 10^5$ cells per reaction. Cells were immobilized onto Concanavalin-A beads (EpiCypher #21-1401) at $5 \times 10^6$ cells/ml and incubated overnight at 4 °C with 0.5 µg antibody against H3K4me1, H3K27ac, H3K4me3, or H3K27me3. pAG-MNase (EpiCypher #15-1016) was added and activated for 2 h at 4 °C. CUT&RUN-enriched DNA was purified using Serapure beads at a 1.4:1 bead-to-DNA ratio. Recovered DNA was quantified by Qubit and normalized to 5 ng input, or the entire reaction was used if <5 ng DNA was recovered.

Libraries were prepared using the CUTANA CUT&RUN Library Prep kit (EpiCypher #14-1001) and sequenced on an Illumina NextSeq 2000 with 2 × 50 bp paired-end reads. Reads were aligned using Bowtie2 v2.2.5 [3]. Multi-aligned reads, duplicates, and ENCODE exclusion-list regions were removed. BigWig files were generated using deepTools v3.5.1 [4], peaks were called using MACS2 v2.2.7.1 [5], and peaks were annotated with HOMER [6].

H3K4me1 and H3K27ac peaks were used to define active/primed chromatin regions in the C/G-transformed cord blood model. H3K4me3 and H3K27me3 CUT&RUN profiles were analyzed in parallel to assess promoter-associated and repressive chromatin redistribution after C/G transduction. Downstream methylation-linked CRE integration focused on H3K4me1/H3K27ac-marked active/primed regions, while H3K4me3/H3K27me3 results are reported as supplementary chromatin redistribution analyses.

*Methyl-enrichment enzymatic methyl sequencing in C/G-transformed cord blood cells*

Day 42 C/G-transformed cord blood cultures were FACS-sorted for the FOLR1+ C/G-associated fraction before methylation profiling. GFP-transduced cord blood controls were processed in parallel. DNA methylation (DNAm) was mapped using CUTANA meCUT&RUN followed by enzymatic methyl sequencing. For each reaction, $1 \times 10^5$ cells were immobilized on Concanavalin-A beads (EpiCypher #21-1401) at $5 \times 10^6$ cells/ml and incubated overnight at 4 °C with MeCP2 (EpiCypher #15-2002). GST antibody (EpiCypher #13-0073) was then added for 30 min at room temperature on a nutator. pAG-MNase (EpiCypher #15-1016) was added and activated for 2 h at 4 °C. Enriched DNA was purified using Serapure beads at a 1.4:1 bead-to-DNA ratio. Recovered DNA was quantified by Qubit and normalized to 5 ng input, or the entire reaction was used if <5 ng DNA was recovered. Libraries were prepared using the NEBNext Enzymatic Methyl-seq v2 kit (NEB #E8015) and sequenced on an Illumina NextSeq 2000 with 2 × 150 bp paired-end reads. Reads were aligned and methylation calls generated using Bismark v0.23.1 [7]. Multi-aligned reads, duplicates, and exclusion-list regions were removed. Differentially methylated regions were identified using methylKit v1.24.0 [8] by comparing C/G-transformed cord blood cells with GFP-transduced controls. DMRs were annotated to promoters, gene bodies, intergenic regions, and cis-regulatory elements and integrated with H3K4me1/H3K27ac CUT&RUN and RNA-seq to identify methylation-linked active/primed regulatory elements.

*C/G occupancy CUT&RUN and leukemia-stage chromatin datasets*

Previously generated M07e C/G CUT&RUN datasets [9] were reprocessed to define leukemia-stage fusion occupancy. CUT&RUN profiles generated with antibodies recognizing the N-terminal CBFA2T3 and C-terminal GLIS2 portions of the C/G fusion were used, and N-terminal GLIS2 CUT&RUN served as a negative-control antibody dataset. Peak files generated in hg19 coordinates were lifted over to hg38 using the UCSC hg19-to-hg38 chain file and CrossMap [10], then sorted and intersected using BEDTools [11]. High-confidence C/G-bound regions were defined as regions shared between N-terminal CBFA2T3 and C-terminal GLIS2 peaks after excluding regions overlapping the N-terminal GLIS2 negative-control peak set. The resulting filtered C/G-bound peak set was used for overlap and enrichment analyses with patient DNAm data. For leukemia-stage chromatin analyses, M07e H3K4me1 and H3K27ac tracks were analyzed in comparison with normal HSC/HSPC controls as indicated. BigWig files were visualized in hg38 coordinates. H3K4me1 and H3K27ac profiles from cord blood and leukemia-stage models were used to define active/primed methylated regulatory regions.

*Integration of C/G occupancy with patient DNAm*

Patient DMCs were intersected with high-confidence C/G-bound regions using BEDTools. DMCs were evaluated globally and after stratification by methylation direction and genomic annotation. For global overlap analysis, the number of hypermethylated or hypomethylated DMCs overlapping C/G-bound regions was counted and expressed as a fraction of total DMCs in each class. For enrichment analysis, Fisher's exact test was performed using all QC-passed EPIC CpGs as the background universe. Odds ratios were calculated for overlap between C/G-bound regions and each DMC class, including promoter, gene-body, and intergenic hypermethylated or hypomethylated CpGs. This analysis identified selective enrichment of C/G-bound regions at promoter-associated hypermethylated CpGs.

*RNA-seq differential expression analysis*

Patient RNA-seq data for C/G+ pAML and NBM controls were obtained from published datasets available through dbGaP (phs000465.v19.p8) [1]. RNA-seq data for C/G–GFP- and GFP-transduced UCB-CD34+ cells were obtained from GSE181726 [2]. Differential expression was analyzed for two primary comparisons: C/G+ pAML versus NBM and C/G-transformed UCB-CD34+ cells versus GFP-transduced UCB-CD34+ controls. DEGs were defined using $|log2FC| \geq 1.5$ and adjusted $P < 0.05$ unless otherwise specified. Patient and UCB DEG lists were intersected to identify patient-only, UCB-only, and shared DEGs. Shared DEGs were classified as concordantly upregulated, concordantly downregulated, or discordant according to the direction of log2FC in the patient and UCB comparisons. Concordantly upregulated genes were used for preranked gene set enrichment analysis and MSigDB Hallmark enrichment as described below.

*Preranked GSEA and MSigDB Hallmark enrichment*

Preranked gene set enrichment analysis (GSEA) was performed to test whether genes concordantly upregulated in both C/G+ pAML and C/G-transformed UCB-CD34+ cells were enriched within patient and UCB ranked transcriptomes. For each comparison, genes were ranked using a signed significance metric: rank metric = sign(log2FC) × −log10(P value). The concordantly upregulated gene set was then tested against the ranked patient and UCB gene lists using preranked GSEA with the standard weighted enrichment statistic [12]. Normalized enrichment score, nominal P value, and false discovery rate were reported where applicable. The same concordantly upregulated gene set was analyzed for MSigDB Hallmark enrichment [13] using an overrepresentation framework, with the tested gene universe defined as genes detected in the corresponding RNA-seq analysis. P values were adjusted using the Benjamini–Hochberg method.

*Definition and visualization of methylation-linked active regulatory elements*

C/G-induced cord blood DMRs were intersected with H3K4me1 and H3K27ac peaks from the C/G-transformed cord blood model to define early active/primed methylated regulatory elements. Hyper-DMRs were classified as active/primed if located within 5 kb of either H3K4me1 or H3K27ac peaks. These regions were linked to nearby genes by genomic proximity and integrated with differential expression to identify methylation-linked transcriptionally permissive CREs. To compare early developmental and leukemia-stage regulatory states, cord blood DMR-expression relationships were integrated with patient EPIC methylation, patient RNA-seq, and M07e leukemia-stage chromatin profiles. Genes were classified as early-specific, early-maintained, or disease-stage-accentuated based on whether methylation-linked regulatory remodeling was detected primarily in the cord blood model, shared between the cord blood model and C/G+ leukemia, or more prominent in patient/M07e leukemia-stage contexts.

Representative loci were selected from the integrated DNAm, chromatin, and expression analyses. Early-maintained loci included *CPZ*, *ITGA2B*, and *MAPK11*. Disease-stage-accentuated loci included *BBC3* and *HPSE2*. H3K4me1 and H3K27ac signal tracks were visualized in hg38 coordinates. In the cord blood model, tracks compared GFP-transduced UCB-CD34+ controls with C/G-transformed UCB-CD34+ cells. In leukemia-stage analyses, tracks compared normal HSC/HSPC controls with M07e cells, with patient hyper-DMCs or DMR-like CpG clusters overlaid where indicated.

Expression log2FC values were summarized for each representative gene using C/G-transformed cord blood versus GFP-transduced cord blood and C/G+ pAML versus NBM comparisons. Differential expression was defined as $|\log2FC| \geq 1.5$ and adjusted $P < 0.05$. Differential methylation was defined as $|\Delta\beta| \geq 0.05$ and adjusted $P < 0.05$ unless otherwise indicated. For genes not detected or not differentially expressed in a given comparison, expression bars were omitted from the corresponding panel.

*HOMER motif enrichment analysis*

Active/primed C/G-CB hyper-DMRs were defined as C/G-induced hypermethylated DMRs located within 5 kb of H3K4me1 or H3K27ac peaks in the C/G-transformed cord blood model. This yielded 1,583 active/primed C/G-CB hyper-DMRs, which were used as the foreground set for known motif enrichment analysis using HOMER against the hg38 genome with input region sizes preserved [6]. Enriched motifs were grouped by transcription factor family for visualization. Because HOMER assigns highly similar ETS-family motifs to multiple ETS factors, ETS motifs were interpreted as ERG/ETS-like motif architecture rather than direct evidence of ERG occupancy. Candidate prioritization was further informed by transcription factor expression in C/G-transformed versus GFP-transduced UCB-CD34+ cells.

*ERG/ETS motif scanning and proximity analysis*

To map ERG/ETS-like motif sites genome-wide, the top ETS-like motif from the HOMER enrichment output was scanned across hg38 using scanMotifGenomeWide.pl. The resulting motif-site BED file was used for proximity analyses and as an IGV-compatible track. For visualization, a uniform-height bedGraph version of the motif-site track was generated to display motif positions as fixed-height vertical marks without altering genomic coordinates. ERG/ETS-like motif proximity to active/primed C/G-CB hyper-DMRs was calculated using bedtools window [11]. Motif proximity was evaluated at cumulative distance thresholds of 250 bp, 1 kb, and 5 kb from each active/primed hyper-DMR. For each threshold, the number and percentage of active/primed hyper-DMRs with at least one ERG/ETS-like motif site within the specified distance were calculated.

*Patient EPIC DMC-to-DMR-like CpG cluster generation*

Patient DNAm data were analyzed from EPIC array profiles comparing C/G+ pAML with NBM controls. Differentially methylated CpGs (DMCs) were defined using an absolute methylation difference ≥5%. CpGs with positive $\Delta\beta$ were classified as hypermethylated in C/G+ pAML, whereas CpGs with negative $\Delta\beta$ were classified as hypomethylated. EPIC CpGs were converted to 1-bp BED intervals in hg38 coordinates. To generate regional methylation units from array-defined DMCs, nearby EPIC DMCs were collapsed into DMR-like CpG clusters. Clustering was performed separately by methylation direction so that hypermethylated and hypomethylated CpGs were not merged. The primary cluster definition used adjacent DMCs within 100 bp with a minimum of two CpGs per cluster. A 200-bp gap definition, also requiring at least two CpGs per cluster, was used as a sensitivity analysis.

*Disease-stage active/primed hyper-DMR-like clusters*

To define disease-stage active/primed methylated regions, patient hyper-DMR-like CpG clusters were integrated with M07e H3K27ac and H3K4me1 peak sets. H3K27ac consensus peaks were generated from three M07e H3K27ac ChIP-seq replicates by retaining regions present in at least two of three replicates. H3K4me1 peaks were generated by sorting and merging the M07e H3K4me1 peak file. Patient hyper-DMR-like clusters were classified as active/primed if located within 5 kb of either M07e H3K27ac or M07e H3K4me1 peaks.

ERG/ETS-like motif proximity to disease-stage active/primed hyper-DMR-like clusters was calculated using bedtools window. Motif proximity was evaluated at cumulative distance thresholds of 250 bp, 1 kb, and 5 kb. The 100-bp DMR-like cluster definition was used as the primary analysis, and the 200-bp definition was overlaid as a sensitivity analysis.

*Locus-level visualization of ERG/ETS-associated regulatory neighborhoods*

Representative loci were visualized in hg38 coordinates using IGV. *ITGA2B* was selected as an early-maintained C/G-linked regulatory target, and *HPSE2* was selected as a disease-stage-accentuated target. Early-stage tracks included C/G-CB hyper-DMRs and C/G-transformed UCB H3K4me1 and H3K27ac signal. Disease-stage tracks included C/G+ pAML hyper-DMCs or DMR-like CpG clusters, M07e H3K4me1 and H3K27ac signal, M07e CBFA2T3 and GLIS2 CUT&RUN signal, M07e ERG ChIP-seq signal, and ERG/ETS-like motif sites. Motif sites were displayed using a uniform-height bedGraph track to preserve genomic position while improving visual interpretation.

*DNMT3B perturbation, rescue, HPSE2 methylation, and expression analysis*

For C/G expression studies, HEK-293T cells were transduced with lentiviral constructs encoding FLAG-GFP control, intact FLAG-tagged CBFA2T3–GLIS2, or FLAG-tagged CBFA2T3–GLIS2 ΔNHR2/ΔZF2-ZF4. The full unmodified FLAG-GFP-HRP-TM_pLX304 plasmid (Addgene #202021) was used as the FLAG-GFP control. FLAG-tagged intact C/G and C/G ΔNHR2/ΔZF2-ZF4 constructs were custom synthesized and sequence-verified by VectorBuilder. FLAG-GFP cells were selected with blasticidin (15 µg/ml), FLAG-C/G cells were enriched by two rounds of sorting for the top 10% brightest GFP-positive cells, and FLAG-C/G ΔNHR2/ΔZF2-ZF4 cells were selected with puromycin (1 μg/ml). Sorted or selected cells were analyzed by immunoblotting. Construct details are provided in Table S8A. Whole-cell lysates were analyzed for FLAG, DNMT1, DNMT3A, DNMT3B, TET1, UHRF1, and β-actin using antibodies listed in Table S8B. TET2 and TET3 antibodies were also tested but did not yield reliable detectable signal under these conditions and were not used for interpretation. Immunoblots are shown as representative experiments.

*DNMT3B*-KO M07e cells were generated by CRISPR/Cas9 targeting of DNMT3B exon 6 as previously described [9]. The sgRNA spacer sequence was 5′-TGCTGTTGCCCGCCGTCTCA-3′. For rescue experiments, DNMT3B-KO cells were transduced with a 3×FLAG-tagged, gRNA-resistant DNMT3B open reading frame containing six synonymous substitutions across the sgRNA target site and a PAM-disrupting mutation, preserving the DNMT3B amino acid sequence while preventing Cas9 re-cutting. DNMT3B depletion and rescue were confirmed by immunoblotting.

Global 5-methylcytosine was measured in M07e WT, DNMT3B-KO, and DNMT3B-rescue cells by LC-MS using stable isotope-labeled internal standards. 5mC and deoxycytidine were quantified from peak-area ratios, and global methylation was reported as percent 5mC relative to deoxycytidine. Three biological replicates were analyzed, and statistical significance was assessed by one-way ANOVA.

Locus-specific DNAm was quantified by pyrosequencing across two HPSE2 upstream promoter-proximal regions in hg38 coordinates: Region 1, chr10:99,235,833-99,236,390; and Region 2, chr10:99,236,772-99,237,316. Region 1 included six quantified CpGs, and Region 2 included four quantified CpGs. Primer sequences, quantified CpG positions, and amplicon information are provided in Table S8C. CpG methylation was compared across M07e WT, DNMT3B-KO, and DNMT3B-rescue cells using three biological replicates. Only successfully quantified CpGs were included in statistical testing.

For HPSE2 expression analysis, RNA was reverse-transcribed using the iScript cDNA Synthesis Kit (Bio-Rad #1708890). HPSE2 expression was measured by RT-qPCR, normalized to ATP5B, and analyzed using the ΔΔCt method. Primer sequences are provided in Table S8D. RT-qPCR data were compared across M07e WT, DNMT3B-KO, and DNMT3B-rescue cells using three biological replicates. Pyrosequencing and RT-qPCR data are shown as mean ± SEM, and statistical significance was assessed by one-way ANOVA.

#### *In vitro drug-response assays*

C/G+ AML cells were plated in 96-well plates at 2 × $10^4$ cells in 100 µl per well. Cells were treated for 48 h with venetoclax (VEN), Nanaomycin-A (Nan-A; DNMT3B-biased inhibitor [14]), azacitidine (AZA), decitabine (DAC), or the indicated combinations. Drugs were serially diluted from 250 µM to 10 nM. DMSO was used as vehicle control at matched concentration. For genetic DNMT3B perturbation, M07e WT and DNMT3B-KO cells were treated with VEN across the dose range. For pharmacologic DNMT3B perturbation, M07e and WSU-AML cells were treated with Nan-A alone or with VEN. For hypomethylating agent (HMA)/VEN combination assays, M07e and WSU-AML cells were treated with AZA, DAC, VEN, or equimolar HMA/VEN combinations.

Cell viability was measured using CellTiter-Glo Luminescent Cell Viability Assay (#G9241, Promega) and normalized to vehicle-treated controls. Experiments were performed in three biological replicates, each with three technical replicates. Dose-response curves were fitted by nonlinear regression in GraphPad Prism, and IC50 values were calculated where appropriate. For HMA priming assays, cells were exposed to AZA or DAC before subsequent VEN challenge. VEN dose-response after HMA priming was compared with matched unprimed VEN controls. AUC values were calculated from replicate dose-response curves and normalized to matched VEN-alone controls for combination assays or unprimed VEN controls for priming assays.

#### *RNA-seq analysis of DNMT3B perturbation and HMA/VEN response*

RNA-seq was analyzed from M07e cells after DNMT3B knockout and after treatment with azacitidine (AZA), venetoclax (VEN), or AZA+VEN, each compared with the matched WT or DMSO control. Differential expression was analyzed using the same pipeline described above, with log2FC and significance thresholds indicated for each analysis. A focused apoptotic-priming heatmap was generated from prespecified genes representing BCL2-family survival, pro-apoptotic priming, and stress-associated apoptotic signaling. The displayed genes were *BCL2*, *BCL2L1*/BCL-xL, *MCL1*, *BBC3*/PUMA, *BAK1*, *DDIT3*/CHOP, and *ATF4*. Significant log2FC values were annotated with asterisks, and non-significant cells were left blank. *PMAIP1*/NOXA was evaluated but excluded because it did not show consistent pro-apoptotic induction across these contrasts.

*TMRM flow cytometry*

Mitochondrial membrane potential was measured using the MitoProbe TMRM Kit for Flow Cytometry (#M20036, Thermo Fisher Scientific). M07e cells were treated for 48 h under matched-dose conditions: DMSO control, AZA 150 nM, VEN 150 nM, or AZA+VEN at 150 nM each. CCCP was included as a depolarization control. Three independent biological replicates were analyzed. After treatment, cells were stained with TMRM at 20 nM for 30 min at 37°C, washed, and analyzed on a BD LSRFortessa 4-laser, 16-color flow cytometer using PE-channel detection. FCS files were analyzed using a reproducible Python workflow. Events were filtered using FSC/SSC parameters, followed by singlet enrichment before TMRM quantification. PE-A was used as the TMRM channel based on separation between DMSO and CCCP-treated controls. TMRM-low thresholds were defined using the midpoint between DMSO and CCCP median log10 fluorescence, with control-percentile thresholds evaluated as sensitivity analysis. Percent TMRM-low cells and median TMRM fluorescence were calculated for each biological replicate.

*Xenograft model and in vivo venetoclax response*

Luciferase-expressing M07e WT and DNMT3B-KO cells were used to assess whether DNMT3B loss enhances venetoclax response in vivo. NSG-SGM3 mice, NOD.Cg-Prkdcscid Il2rgtm1Wjl Tg(CMV-IL3,CSF2,KITLG)1Eav/MloySzJ, also known as NSGS,

NSG-SGM3, or NSG-Triples (JAX #013062; RRID: IMSR_JAX:013062), were injected by tail vein with 0.25 x 10^6 M07e WT or DNMT3B-KO cells per animal. Six mice were included per group, with three males and three females. After a 7-day engraftment period, mice were treated with venetoclax at 25 mg/kg by oral gavage, 5 days/week, from week 2 through week 9.

For bioluminescent imaging (BLI), mice were injected intraperitoneally with D-luciferin at 150 mg/kg and imaged using an IVIS imager every 7 days. D-luciferin was prepared at 15 mg/ml in sterile DPBS, and 200 µl was injected per 20 g mouse before imaging. Venetoclax was prepared at 2.5 mg/ml in 15% Cremophor EL/85% saline, and 200 µl was administered per 20 g mouse by oral gavage. Leukemia progression was monitored longitudinally by BLI. Where collected, endpoint leukemia burden was assessed by spleen involvement and flow cytometric quantification of FOLR1-positive leukemic cells in bone marrow. Mice were monitored for humane endpoint criteria, including reduced activity, sluggishness, hind-limb paralysis, or other protocol-defined morbidity. Overall survival was calculated from cell injection and compared by log-rank test. Animal studies were performed under an approved institutional IACUC protocol.

*Statistical analysis*

Statistical analyses were performed using GraphPad Prism, R, and Python. Comparisons between two groups were performed using two-sided t-tests unless otherwise indicated. Comparisons among three or more groups were performed using one-way ANOVA unless otherwise indicated. Multiple testing was corrected using false discovery rate procedures where appropriate. Fisher's exact test was used for categorical overlap and enrichment analyses, including overlap between C/G-bound regions and methylation classes. Kaplan-Meier survival curves were compared using log-rank tests. Dose-response curves were fitted by nonlinear regression, and IC50 and AUC values were calculated from replicate dose-response curves. Data are shown as mean ± SD or mean ± SEM, as indicated in the corresponding figure legends. Statistical tests were applied to independent biological replicates unless explicitly stated.

*Software and reproducibility*

Genomic interval processing was performed using BEDTools, CrossMap, and custom Python/R scripts. Sequencing reads were aligned using Bowtie2 or Bismark, as appropriate. BigWig files were generated using deepTools, peaks were called using MACS2, and motif analyses were performed using HOMER. Differential methylation was analyzed using methylKit. Genomic analyses were performed in hg38 coordinates unless otherwise stated. CUT&RUN, ChIP-seq, and methylation signal tracks were visualized in IGV. Source data used for figure generation were exported as tab-delimited or Excel files. Code used for figure generation and reproducible analysis will be deposited on GitHub upon publication.

### Supplementary Results

#### C/G transduction redistributes promoter-associated and repressive chromatin marks

To determine whether C/G-induced chromatin remodeling extended beyond active/primed enhancer-associated marks, we also examined H3K4me3 and H3K27me3 CUT&RUN profiles in the UCB model. H3K4me3 showed strong peak-centered enrichment in GFP-control UCB-HSPCs that was markedly attenuated in C/G-transformed cells. Among 16,414 H3K4me3-centered regions, 251 regions showed newly detectable signal, 111 baseline-positive regions were amplified, and 15,836 were decreased, corresponding to a median fold-change of 0.14. In contrast, H3K27me3 showed focal gain in C/G-transformed cells. Among 4,461 H3K27me3-centered regions, 344 regions showed newly detectable signal and 3,173 baseline-positive regions were amplified, with a median fold-change of 3.59. These data indicate that C/G transduction causes broader chromatin reorganization; however, because the strongest methylation-linked CRE signal in the main analysis was associated with H3K4me1/H3K27ac-marked active/primed regulatory regions, downstream integration focused on these enhancer-associated marks (Supplementary Fig. S1).

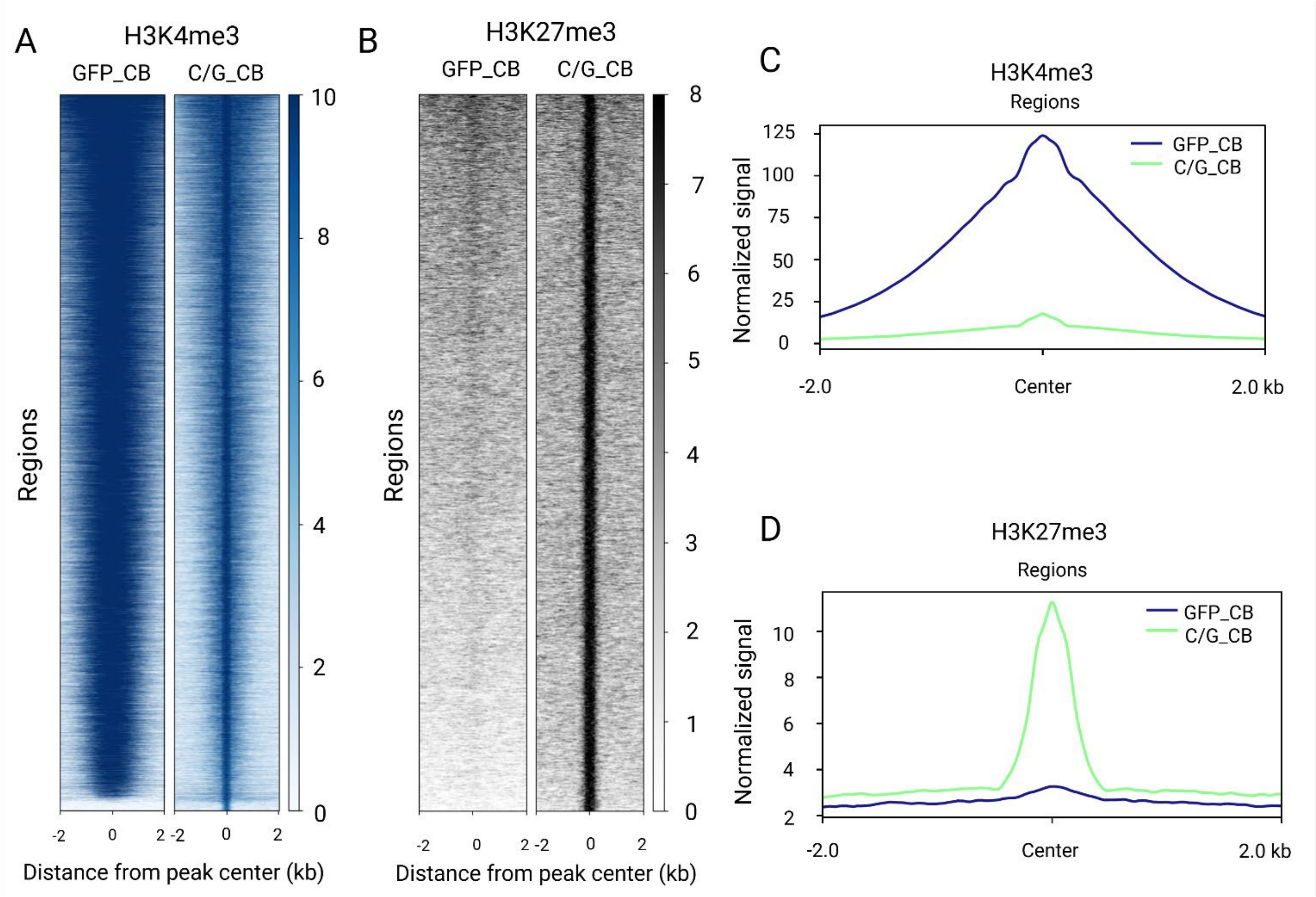


**Figure S1. C/G-associated redistribution of H3K4me3 and H3K27me3 in the UCB model.**

(**A,B**) Heatmaps of H3K4me3 and H3K27me3 CUT&RUN signal centered on peak regions in GFP-control and C/G-transformed UCB-HSPCs.

(**C,D**) Aggregate H3K4me3 and H3K27me3 signal profiles across the same peak-centered regions. Signal is shown across ±2 kb from peak centers. H3K4me3 showed broad attenuation after C/G transduction, whereas H3K27me3 showed focal enrichment in C/G-transformed cells.
